# Mutation-Attention (MuAt): deep representation learning of somatic mutations for tumour typing and subtyping

**DOI:** 10.1101/2022.03.15.483816

**Authors:** Prima Sanjaya, Sebastian M. Waszak, Oliver Stegle, Jan O. Korbel, Esa Pitkänen

## Abstract

Cancer genome sequencing enables accurate classification of tumours and tumour sub-types. However, prediction performance is still limited using exome-only sequencing and for tumor types with low somatic mutation burden such as many pediatric tumours. Moreover, the ability to leverage deep representation learning in discovery of tumour entities remains unknown. We introduce here Mutation-Attention (MuAt), a deep neural network to learn representations of simple and complex somatic alterations for prediction of tumour types and subtypes. MuAt achieved prediction accuracy of 89% for whole genomes (24 tumour types) and 64% for whole exomes (20 types), and a top-5 accuracy of 97% and 90%, respectively. Tumour representations learnt by MuAt included tumour entities such as acral melanoma, SHH-activated medulloblastoma, *SPOP*-associated prostate cancer, microsatellite instability, and *MUTYH*-associated pancreatic endocrine tumours although these tumour subtypes and subgroups were not used as training labels. Integrated representations of somatic alterations hold significant potential to drive discovery of novel tumour entities and clinical application.

## 1 Introduction

Accurate identification of tumour histological type and molecular subtype is crucial to determining cancer diagnosis, prognosis and treatment choice [1, 2]. In pediatric brain tumors, long-term survival can range from 90% for WNT-medulloblastomas to 40% for Group 3-medulloblastomas [3]. Solid tumours exhibiting microsatellite instability (MSI) resulting from defective mismatch repair (MMR) are susceptible to treatment with PD-1 immune checkpoint inhibitors leading to improved response and survival rates [4, 5]. Moreover, approximately 3-5% of metastatic cancers do not have a clear primary site of origin despite comprehensive clinical workup [6, 7]. These cases, termed cancers of unknown primary (CUPs), present a challenge as targeted treatment options depend on the tissue of origin. CUPs are thus often treated with broad spectrum antineoplastic drugs with limited success, instead of site-specific treatments. Liquid biopsies can be used to detect circulating tumor DNA (ctDNA) originating from cancer cells before metastatic spread and to predict disease outcome [8, 9, 10]. Similarly to CUPs, determining the tissue of origin of ctDNA is a key obstacle in enabling clinical action.

Somatic mutations in a cancer cell are the consequence of the mutational processes which acted on its ancestors in the somatic cell tree [11]. Many such processes have been identified, including exogenous processes such as ultraviolet radiation and polycyclic aromatic hydrocarbons in tobacco smoke, and endogenous processes such as spontaneous deamination of methylated cytosines, defective DNA repair, and DNA replication infidelity [12]. These processes can have distinct characteristics in terms of DNA substrate preference (*e*.*g*., CpG, mononucleotide microsatellite), mutation type (*e*.*g*., single-base substitution, insertion or deletion, or structural alteration) and genomic position (*e*.*g*., intronic or late replicating region preference), among others [13]. Typically only a handful of mutational processes are active in a cell of specific type and location within the body and tissue [14, 15]. For instance, skin cells exposed to the sun are susceptible to DNA damage due to ultraviolet radiation, whereas B cells undergo somatic hypermutation affecting predominantly the immunoglobulin heavy chain variable region of the genome. Somatic mutations can thus be informative of the tissues and conditions where the mutations occurred, and consequently, cancer genome sequencing can be used to scrutinize the somatic mutations of a cancer with the prospect of revealing its tissue of origin and molecular subtype.

Recently, several computational methods have been developed to predict tumour types by analyzing somatic driver and passenger mutation patterns in next-generation sequencing data [16, 17]. TumorTracer is a random forest classifier combining copy number profiles and nucleotide substitution spectra attaining 85% and 69% accuracy across 6 and 10 primary sites, respectively [18]. Soh *et al*. predicted tumour types with a support vector machine using information on somatically mutated genes resulting in 49% accuracy in 28 tumour types, with addition of copy number profiles increasing accuracy to 78% [19]. Especially in whole cancer genomes, the mutational landscape is dominated by passenger mutations which are highly informative of the tissue-of-origin. As part of the Pan-Cancer Analysis of Whole Genomes project (PCAWG), Jiao *et al*. explored tumour type prediction in 2,606 tumours representing 24 tumour types, and 88% accuracy in an independent set of tumour whole genomes with a deep neural network model which takes as input counts of mutation types and their binned genomic positions in each tumour [20]. Tumours exhibiting microsatellite instability (MSI) were removed from data prior to model training. Both Jiao *et al*. and Salvadores *et al*. [21] found the utility of driver mutations in accurately predicting tumour types to be limited due to the relatively small number of driver alterations per tumour, few recurrent driver alterations, and lack of strong tumour type specificity for cancer driver genes. Recently, Danyi *et al*. showed data augmentation to be an effective strategy for tumour typing with sparse sequencing data such as sequencing of ctDNA [22].

While supervised approaches have been developed to predict tumour subtypes [23, 24], unsupervised methods are more common due to lack of labeled subtype data. In unsupervised tumour subtyping, one typically aims to find a compact set of latent factors explaining the observed data, often compassing multiple modalities [25], and then identifying subtypes using latent factors. Recent subtyping methods have employed matrix factorization [26], clustering [27], deep autoencoders [24], and adversarial learning [28] of multiomics data. Discovery of prognostic subtypes has been done by weighting or selecting features based on survival [24, 29]. Sequence context of somatic mutations has been shown to be informative in subtyping of breast cancers [30].

Here, we developed a novel deep neural network (DNN) model, termed Mutation-Attention (MuAt), which allows us to predict tumour types from cancer whole-genome and whole-exome sequencing data. It leverages the ability of DNNs to work in a supervised setting to learn representations that can be used to explore and explain the structure of input data beyond class labels. MuAt utilizes a recent innovation in deep learning called the attention mechanism [31, 32]. This mechanism allows deep neural networks to focus on data elements which are important to solving the learning task at hand, often leading to improved model performance and explainability. MuAt is able to integrate single-nucleotide and multi-nucleotide substitutions (SNVs/MNVs), short insertions and deletions (indels), structural variant (SV) breakpoints, and combinations of these primary genetic alterations. In contrast to previous approaches, MuAt integrates mutation type and genomic position information at a per-mutation level, instead of representing type and positions as aggregated counts.

Our models achieve high accuracy in predicting tumour types, with top-1 and top-3 accuracies of 88.9% and 96.1% in the 24 tumour types that were studied within the PCAWG consortium. It further outperforms the previous state-of-the-art approaches for cancer types that have been challenging to predict such as tumours with MSI. We investigate the utility of our model in tumour exome sequencing data from the TCGA consortium, achieving 64.1% accuracy across 20 tumour types. Exploring the representations learnt by MuAt, we show that the model learns to differentiate tumour subtypes which were not given as input information. These subtypes include tumours driven by somatic and germline mutations such prostate cancers with somatic *SPOP* mutations and pancreatic endocrine tumours with germline *MUTYH* mutations, hypermutable subtypes such as microsatellite-unstable cancers and polymerase *ϵ* proofreading deficient tumours, as well as *CCND1*-amplified acral melanomas, and Sonic Hedgehog (SHH)activated medulloblastomas.

The use of attention mechanism together with the ability to learn representations for different data modalities such as mutation types and positions allows MuAt to represent each mutation as a combination of these modalities. To gain insight into model results, we show that the trained model learns to focus its attention to mutations that are characteristic for each tumour type. MuAt models trained with cancer genomes from PCAWG and TCGA consortiums, an interactive browser, and the source code are available under a permissible license at GitHub (https://github.com/primasanjaya/mutation-attention).

## 2 Data

To train MuAt models, we utilized WGS data from the Pan-Cancer Analysis of Whole Genome (PCAWG) project and WES data from the Pan-Cancer Atlas project of TCGA. PCAWG analyzed whole genomes of 2,658 human tumours and matched normal samples across 38 tumour types obtained from International Cancer Genome Consortium (ICGC) and The Cancer Genome Atlas (TCGA) donors [33]. The project released a dataset of somatic mutations called uniformly in these tumours containing somatic SNVs, MNVs, indels (<50 bp), SVs and mobile element insertions (MEIs). To train MuAt models, we utilized only tumour types with more than 20 tumours in PCAWG resulting in 2,595 tumours across 24 tumour types and 18 primary sites. These tumours harbored a total of 47,649,187 somatic mutations, divided into 41,969,899 SNVs, 826,093 MNVs, 3,720,396 indels, 1,109,524 SVs and 16,757 MEIs, constituting the PCAWG training dataset.

The Pan-Cancer Atlas project of TCGA [34] released the MC3 somatic mutation dataset consisting of a total of 5,717,732 somatic mutations from 8,942 tumours across 32 tumour types [35]. We selected the 20 tumour types with more than 100 tumours into our TCGA training dataset, resulting in 7,352 tumours and 2,682,344 somatic mutations (2,498,826 SNVs, 46,589 MNVs, and 136,929 indels). No SVs in the data were included in the training dataset due to these events often occurring in the intergenic regions and thus not adequately captured by exome sequencing.

To validate our results with data which was not used in training, we used the whole genomes available in ICGC that were not included in the PCAWG dataset above. Out of 16 tumour types, 5 tumour types were matched to PCAWG tumour types, while the rest were used to evaluate the behaviour of MuAt when predicting unseen tumour types. There were 1,072 tumours, having in total 12,099,773 mutations (8,180,922 SNVs, 233,535 MNVs and 3,685,316 indels). SVs were not available in this dataset. Further details on the datasets can be found in Supplementary Table 1.

## 3 Model

MuAt is a DNN model, which predicts tumour types based on a catalogue of somatic alterations that are observed in a single cancer genome (Figure 1). We describe here briefly the key aspects of MuAt, and provide details in **Methods** and Supplementary Figure 1. The model consists of three consecutive modules. In the first module, mutations are encoded and embedded into a feature space. Three sources of information are used to encode each mutation: 1) mutation type embedded in a three-nucleotide sequence context (*e*.*g*., Ap[C>T]pG, Tp[delC]pC), 2) genomic position in 1-Mbp bins, and 3) annotations describing whether a mutation occurs in a gene or in an exon, and the coding strand orientation. The supported somatic mutation types are SNVs, indels and SV breakpoints. The MuAt encoding allows for combinations of up to three of these simple mutations to be represented in the sequence context, for instance MNVs (*e*.*g*., Cp[C>T]p[C>T]) or >1 bp indels (*e*.*g*., [insT]p[insT]p[insT]). Sequence contexts, genomic positions and annotations are represented as one-hot encoded vectors. MuAt learns feature embeddings of these three modalities, which are then concatenated and used as input to the second module.

**Figure 1:**
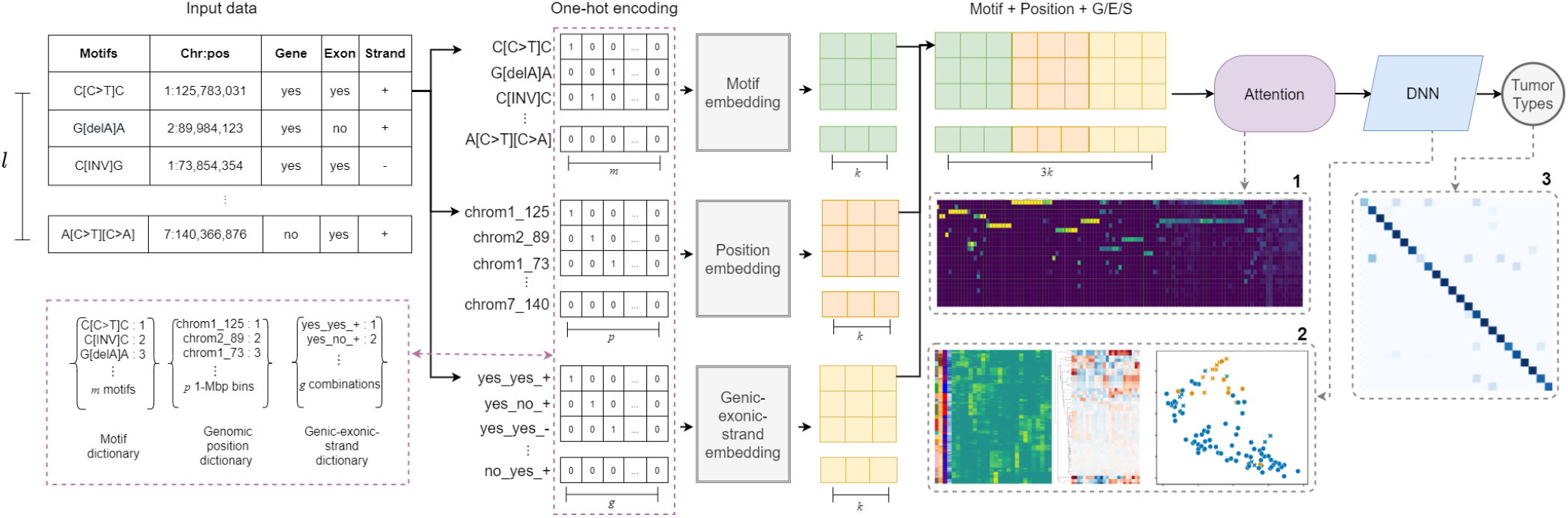
Illustration of the MuAt deep neural network to predict the type of a tumour from its catalogue of somatic mutations. First, mutation data is one-hot encoded. MuAt integrates three data modalities: 3-bp sequence motif, genomic position and genomic annotations. Then, embedded mutation vectors are fed to the attention mechanism. Finally, mutation-level features are combined into tumour-level features, and tumour type is predicted. MuAt models can be interrogated by analysing 1) the attention matrix to recover informative mutations for tumours and tumour types, 2) tumour-level features for tumour subtype discovery, and 3) prediction performance.

In the second module, an attention mechanism is used to assign more weight (“soft-select”) to pairs of mutations which are informative in predicting the tumor type, and compute input features for the third module. Attention is defined as

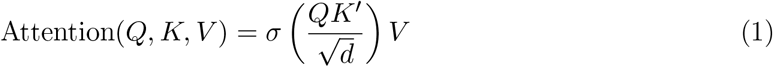

where *Q, K, V* are called the query, key and value matrices, respectively, and *σ* denotes the softmax [31, 32]. *Q, K* and *V* are all *l* × *d* matrices, where *l* is the number of mutations and *d* is the feature dimension; these matrices are obtained as linear transformations of the input. Product *QK*′ can be seen as a similarity matrix, with the softmax being used to soft-select the most relevant mutations (“keys”) for each mutation (“query”). The attention thus maps each input mutation to a *d*-dimensional feature space defined in terms of similarity to other mutations in the same tumour. Unlike many natural language processing applications which employ attention [32], no order is imposed on the mutations: MuAt treats somatic mutations analogously to a bag-of-words.

The third module combines the mutation features with fully connected layers yielding tumor-level features. These features are used to compute the final output of the model, which are probabilities over the tumor type labels. The three modules constitute a single model, where all parameters are trained end-to-end with backpropagation and stochastic gradient descent. The trained MuAt model can be interrogated by extracting mutation-level features from the attention module and tumor-level features from the last module. We can project the latter onto a two-dimensional space with UMAP [36] for discovery of tumour subtypes.

## 4 Results

### Evaluation of histological tumour typing performance

We first evaluated the contribution of different somatic mutation types and mutation annotations to cross-validated prediction performance. In tumour whole genomes from the PCAWG consortium, the best MuAt performance was obtained with the combination of SNVs, MNVs, indels and genomic position (accuracy 88.8%, 97.6% top-5) (Fig. 2a), although the differences in performance between these scenarios tested were relatively small. In tumour exomes from the TCGA consortium, addition of indels and mutation annotations improved the performance substantially over the other WES models (accuracy 64.1%) (Fig. 2a & Supplementary Fig. 3). While predicting the exact tumor type correctly with somatic mutations from exomes compared to whole genomes was more challenging, MuAt reached a high top-5 accuracy of 90.6%. Furthermore, MuAt predictions were found to be reliable, indicating that the model was well-calibrated (Supplementary Fig. 4).

**Figure 2:**
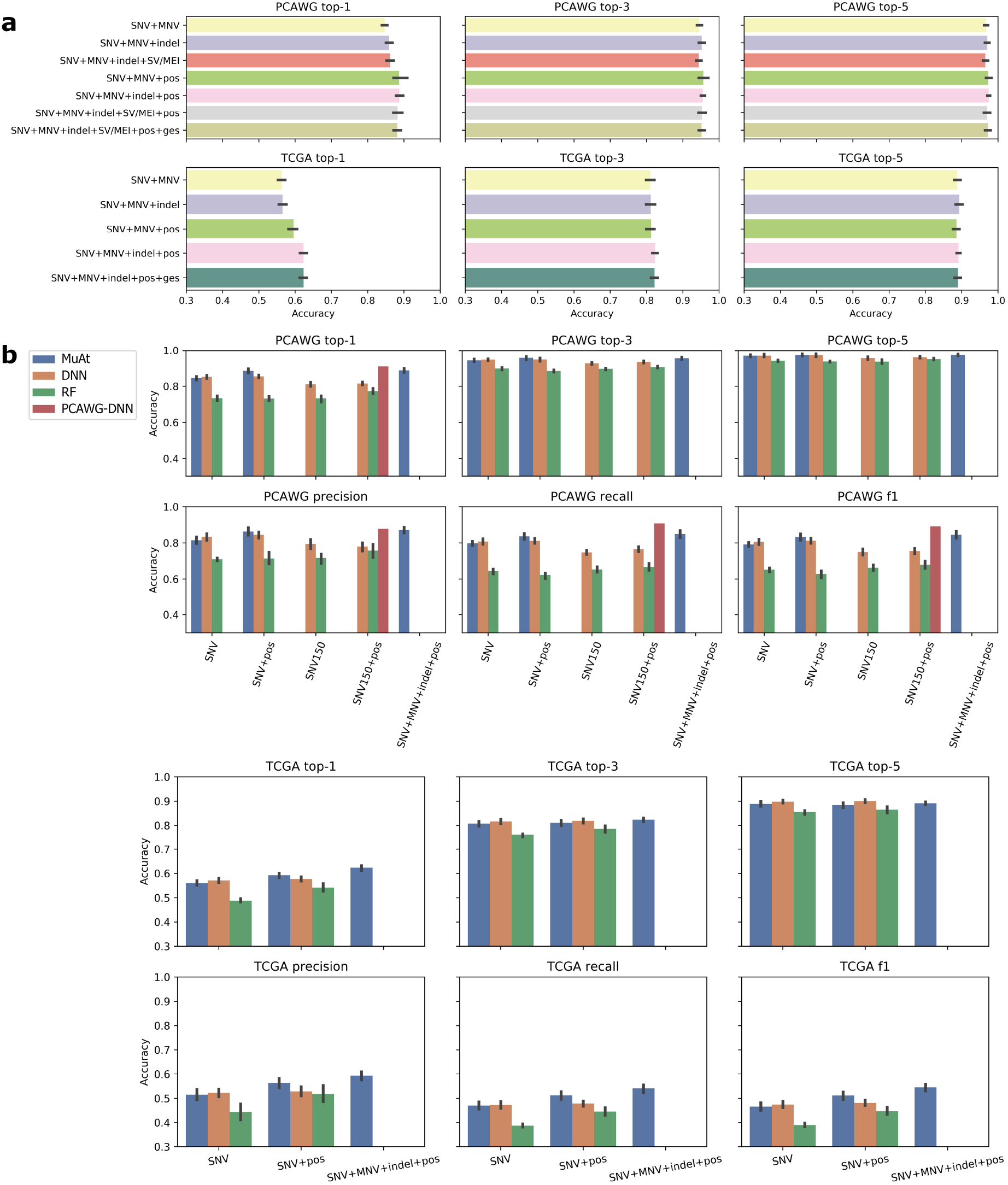
**a**) Top-1, top-3 and top-5 accuracies of MuAt models trained with different mutation types in PCAWG cancer genomes (top) and TCGA cancer exomes (bottom). Cross-validation accuracies are shown, with the standard deviation of accuracy in cross-validation folds indicated by error bars. **b**) Benchmarking MuAt prediction performance in PCAWG genomes and TCGA exomes against a recent DNN model [20] and a random forest model (RF). Two versions of the DNN model performance are reported, analysis of the PCAWG dataset (DNN) and the performance reported in [20] (PCAWG-DNN). SNV150 indicates the feature set used in [20], “ges” stands for genic, exonic and strand attributes (**Methods**). The last column in the PCAWG comparison shows the best performance obtained by MuAt (SNV+MNV+indel+position). Note that the data range starts from 0.3 in all charts.

We then benchmarked our method against the current state-of-the-art. Here, MuAt outperformed the recent DNN model of Jiao *et al*. [20], which achieved 85.5% accuracy with PCAWG cancer genomes and 59.4% accuracy in TCGA cancer exomes (Fig. 2b). In [20], the method was evaluated in PCAWG data where MSI tumours had been removed. As the exact type of an MSI tumour can be difficult to predict, and since they represent a clinically important subgroup [37], we kept the MSI tumours in the data. This may explain the difference between our experiment and their reported accuracy of 91% [20]. The accuracy of a random forest model was substantially worse than either model (77.3% PCAWG, 57.7% TCGA). Full results are available in Supplementary Table 2 and Supplementary Table 3.

For the remainder of experiments, we proceeded with the MuAt model trained on SNVs, MNVs, indels and SVs/MEIs, and genomic positions in PCAWG data (**Methods**). We first investigated how the number of mutations in each tumour influences prediction performance. As expected, tumours with smallest mutational burden (*n*=259 tumours, <1,109 mutations) showed the poorest prediction accuracy with 78.4% of tumours correctly predicted (Fig. 3a). Many prostate and thyroid cancers, and medulloblastomas with low burden were found hard to predict, whereas pilocytic astrocytomas were predicted more accurately. Similarly, the tumours with the highest burden (*n*=256 tumours, >29,626 mutations) were more difficult to predict (accuracy 82.8%) than the tumours with intermediate burdens (1,109–29,626; accuracy 90.8%). This group included many colorectal, stomach and uterine cancers with DNA repair defects leading to hypermutability. In general, the model misclassified similar tumour types, or tumour types exhibiting similar mutational mechanisms, such as lung cancers and gastrointestinal cancers (Fig. 3b). Another reason for tumour misclassification might be the capacity of the MuAt model being limited to 5,000 mutations that are randomly sampled from the mutation catalogue of the tumour (**Methods**). This capacity was exceeded in tumours with highest mutation counts (Fig. 3a).

**Figure 3:**
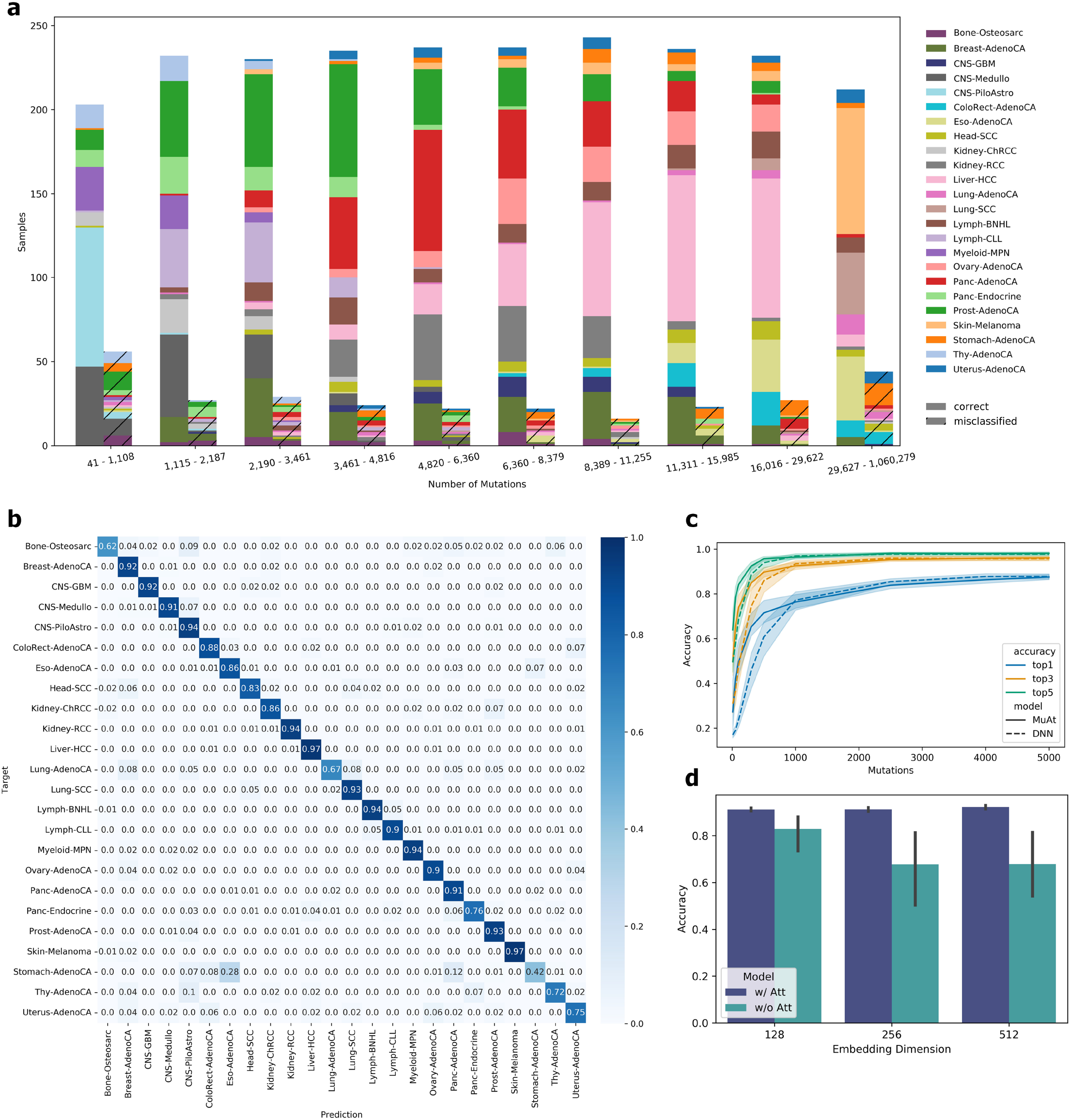
**a**) MuAt predictions in PCAWG cancer genomes stratified by the number of somatic mutations and whether the tumour type was correctly (solid colours) or incorrectly (crosshatched colours) predicted (top-1). Cross-validation results reported for PCAWG dataset. **b**) Confusion matrix of the best-performing MuAt model in PCAWG data. **c**) Comparison of MuAt and DNN [20] accuracy (Y-axis) on sparse data with respect to the number of mutations in downsampled tumours (X-axis). Top-1/3/5 accuracies are shown. **d**) Accuracy of MuAt with attention (w/ Att) and without (w/o Att), and with respect to the embedding dimensionality.

We investigated the performance of the MuAt PCAWG model in sparse data by subsampling mutations. Figure 3c shows the knee in accuracy (71.1%) to occur already at around maximum number of 500 mutations per sample, and steadily increasing up to 5,000 mutations. Notably, top-5 accuracy with 500 mutations was relatively high (95.6%). MuAt performed slightly better than the DNN model in subsampled data. To understand the contribution of the attention mechanism, we evaluated MuAt models with the attention module removed. Such models were found to train and perform poorly compared to full MuAt model (Fig. 3d, Supplementary Fig. 5).

Finally, we evaluated MuAt with data not used in training to assess the method’s ability to generalize to new data. To do this, we first constructed a MuAt ensemble model from the ten best models found in cross-validation (**Methods**). We then predicted tumour types in the subset of ICGC whole genomes which were not part of PCAWG and where tumour types matched those in the PCAWG (six cohorts, five tumour types). Only somatic SNVs and MNVs were used to train the ensemble model, since these were the only mutation type which was available for all six cohorts. In addition, genomic positions and mutation annotations were used. The model achieved 84.8% mean accuracy across all tumours, with the individual cohorts predicted at 74- 100% accuracy (Supplementary Fig. 7). While in training data medulloblastomas were predicted at 91% accuracy, in an independent pediatric medulloblastoma cohort (PEME-CA) the MuAt ensemble model reached only 74% top-1 accuracy. Nevertheless, MuAt predicted tumours of this cohort to be one of the three central nervous system tumour types present in the training dataset in 96% of cases.

### MuAt distinguishes molecular tumour subtypes

We explored the MuAt tumour-level features (n=24, M_1_,…,M_24_) learnt in PCAWG data by projecting the features to a two-dimensional space with UMAP [36]. PCAWG tumours clustered by tumor type as expected due to the high performance of the model in predicting tumor types (Fig. 4). We then investigated whether MuAt features learnt by classifying tumor types could be informative of histological or molecular subtypes even though information on subtypes was not provided during model training. By correlating MuAt features with known or predicted driver events reported in PCAWG tumours [33] and correcting for tumour histology (**Methods**), we identified a striking association of MuAt features with somatic driver events in *SPOP* (*q*=6.591×10^−12^; Supplementary Fig. 8), a candidate driver gene in prostate cancer [38]. All twelve prostate cancers with *SPOP* mutations clustered in the MuAt feature UMAP (Fig. 4a). These tumours harbored 2.3 times (95% CI, 1.2–4.3x) more somatic SVs than wildtype tumours (Supplementary Table 4). In contrast to tumours with *BRCA1* or *BRCA2* driver events, which harbored 1.6x (95% CI, 1.0–2.4x) and 1.4x (95% CI, 0.9–1.9x) more SVs as well as excess of 10– 100 bp deletions (*BRCA1*, 2.2x; *BRCA2*, 7.3x), *SPOP* tumours did not display excess of other somatic mutation types. Instead, *SPOP* tumours constituted a molecular subgroup of prostate cancer characterized by increased somatic structural alteration burden, compatible with previous reports [39]. Prostate cancers with *ERG* driver mutations (*n*=85), mutually exclusive to *SPOP* (OR=0.102, 95% CI, 0.002–0.725, *p*=0.013; [40]), were evenly distributed among the remaining prostate cancers in MuAt clustering and somatic translocation (Supplementary Fig. 9).

**Figure 4:**
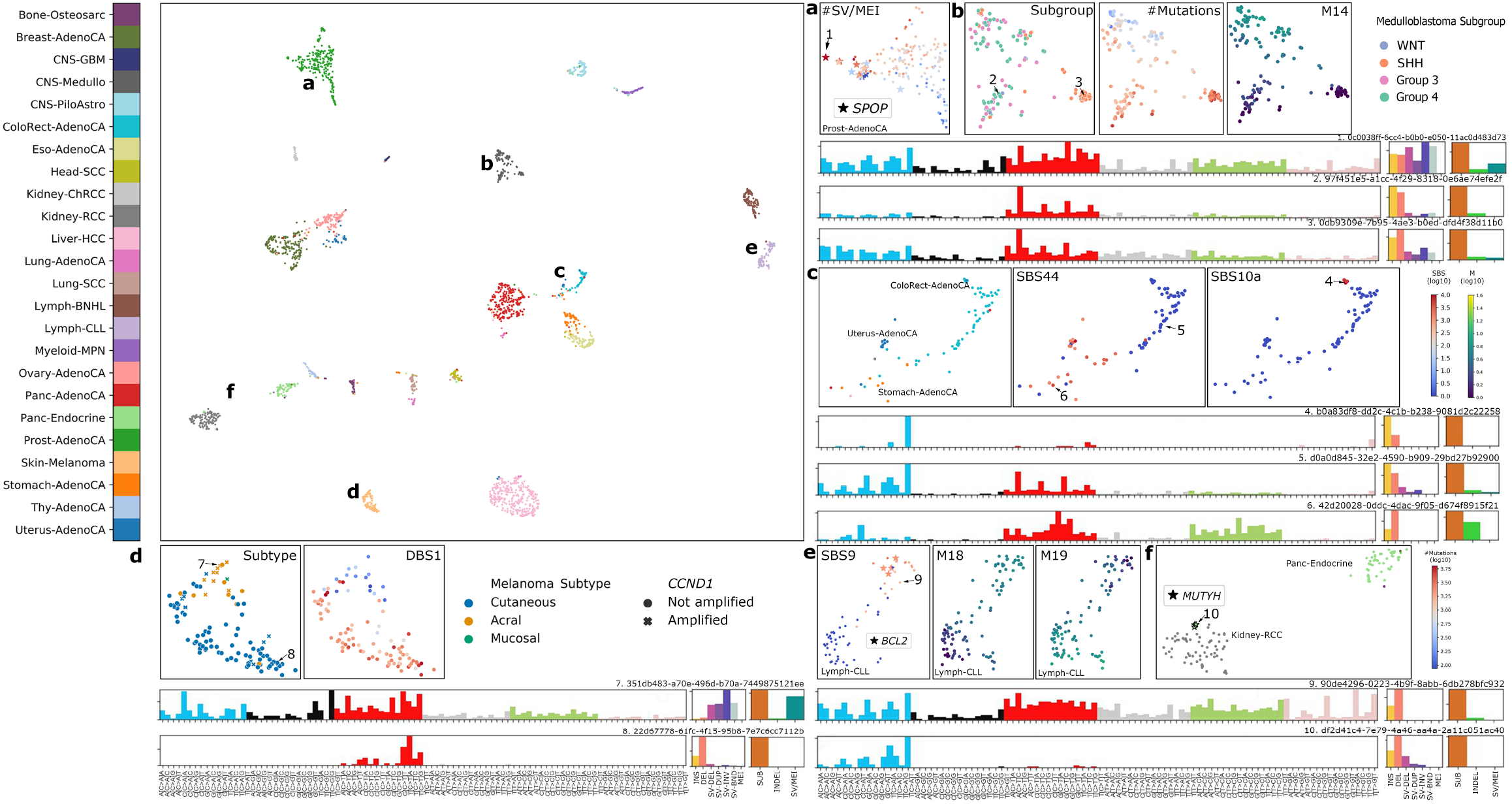
UMAP projection of MuAt tumour-level features in PCAWG data. MuAt recognized tumour subtypes in **a**) prostate cancers with a subgroup defined by *SPOP* driver mutations and increased somatic structural variant burden, **b**) medulloblastomas, **c**) colorectal cancers with microsatellite-unstable (MSI), polymerase *ϵ* deficient, and microsatellite-stable (MSS) cancers, **d**) skin melanomas subtypes with *CCND1* amplifications, **e**) chronic lymphocytic leukemias associated with somatic hypermutability, and **f**) pancreatic endocrines with germline *MUTYH* mutations. Specific example tumours are indicated by numbers, with SNV, indel and SV types, and SNV/indel/SV proportions shown.

In medulloblastomas, four subgroups were visible in the feature UMAP (Fig. 4b). One of these was found to correspond to the Sonic Hedgehog (SHH)-activated medulloblastomas of adult patients with mutation landscapes dominated by the age-associated CpG>TpG substitutions [41]. Furthermore, *PTCH1, DDX3X*, and *SMO* driver mutations characteristic to SHH medulloblastomas, and *PRDM6* enhancer hijacking driver events found in Group 4-medulloblastomas associated with MuAt features (FDR<1%; Supplementary Fig. 8). Interestingly, medulloblastomas with *PRDM6* driver alterations (*n*=7) were confined to one of MuAt feature clusters (Supplementary Fig. 10. These tumours displayed an increased genome-wide burden of SV duplications (3.7x, 95% CI, 1.6–8.3x), inversions (2.4x, 1.0–5.4x) and translocations (2.4x, 1.0–5.6x), but no excess of other mutation types compared to wildtype medulloblastomas (**Methods**).

MuAt identified tumours with mismatch repair deficiency (MMR) resulting in microsatellite instability as well as tumours exhibiting very high burden of mutations, especially TpCpT>TpApT substitutions characteristic of polymerase *ϵ* and *δ* proofreading deficient tumours (Fig. 4c). Driver mutations in the MMR gene *PMS2* associated with MuAt feature M_6_ (*q*=5.53×10^−7^). The fraction of somatically mutated microsatellites, a measure for the level of MSI in a tumour, positively associated with features M_6_, M_9_ and M_13_ (*q <*0.05; Supplementary Table 5; **Methods**). TpCpT>TpApT substitutions positively associated with 11 MuAt features at 1% FDR, with *M*_3_ displaying the strongest association (Supplementary Table 6).

In skin melanomas, we observed acral melanomas to cluster by MuAt features (Fig. 4d). Tumours in this subgroup displayed many somatic SVs and amplifications of *CCND1*, a common alteration in acral melanomas associating with ulceration and localized metastasis [42]. The only mucosal melanoma in the data harbored a *CCND1* amplification and clustered with the acral melanomas. The remaining melanomas, mostly of the cutaneous subtype, had a high number of CpC>TpT dinucleotide substitutions compatible with signature DBS1 due to UV light exposure. Finally, in chronic lymphocytic leukemias, MuAt differentiated tumours with patterns of somatic mutations that had occurred in B cells during *IGH* gene rearrangements (Fig. 4e) [43].

Pancreatic neuroendocrine tumours (PanNETs) of four patients clustered unexpectedly with kidney cancers (Fig. 4f, Supplementary Fig. 11). Accordingly, three of these four PanNETs were misclassified by MuAt as kidney cancers. These patients were found to be carriers of germline mutations in *MUTYH* (p.Tyr176Cys, two patients; p.Pro292Leu; c.924+3A>C). (Supplementary Table 7), and all four tumours showed loss-of-heterozygosity of *MUTYH. MUTYH* encodes a DNA glycosylase involved in base excision repair, and germline *MUTYH* mutations have been implicated in a specific G:C>T:A somatic mutation signature found in PanNETs and colorectal cancers [44, 45]. Consistently with these earlier results, we saw an excess of C>A substitutions in NpCpA and NpCpT contexts in *MUTYH* tumours compared to other PanNETs in PCAWG (*t*=9.63, *p*=6.57×10^−15^) (Supplementary Fig. 11).

### MuAt features are informative of mutational patterns and correlate with mutational signatures

A subset of histologies in PCAWG were characterized by high values of specific features (Fig. 5a, Supplementary Fig. 12) such as central nervous system tumours by feature M_1_, lung cancers by M_3_, pilocytic astrocytomas by M_4_, gastrointestinal tract cancers by M_6_, and liver cancers by M_9_. Four features associated with higher (M_6_, M_8_) or lower (M_4_) overall somatic SNV burden (Supplementary Fig. 16, Supplementary Table 6). Other features associated with more specific mutational patterns, such as higher structural variant counts (M_15_, M_18_, M_22_), 1-3 bp indels (M_3_, M_6_, M_8_, M_13_, M_21_, M_24_), large deletions and duplications (M_5_), or only duplications (M_1_). Specific SNV triplet patterns included TpCpA>TpApA and TpCpT>TpApT (M_20_), and TpCpN>TpGpN and TpCpN>TpApN patterns matching the mutational footprint of APOBEC activity (M_15_, M_20_).

**Figure 5:**
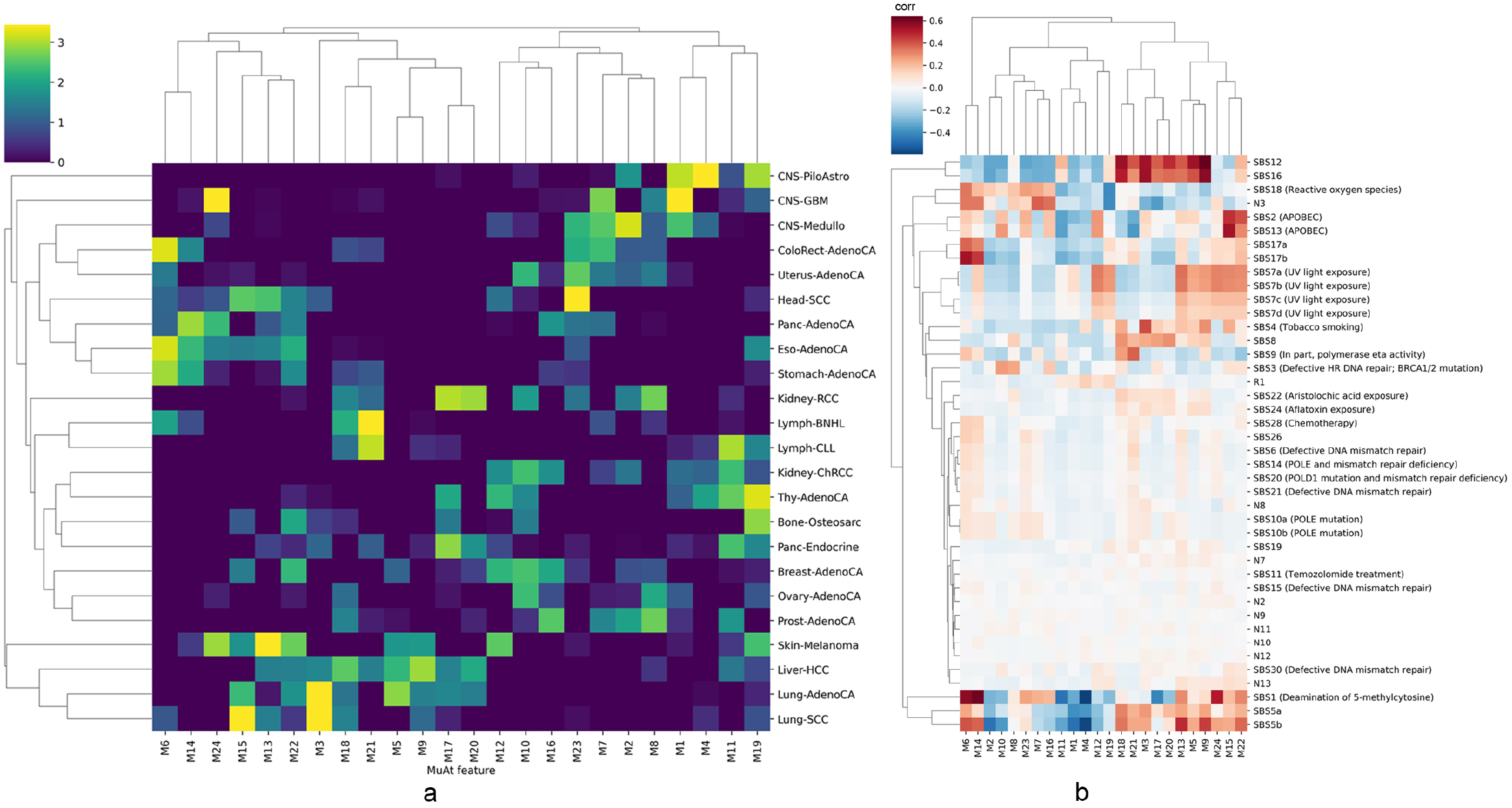
Association of MuAt tumour-level features with tumour types and mutational signatures. **a**) MuAt mean feature values in PCAWG genomes. Tumour type (histology) is indicated on the right. **b**) Rank correlation between MuAt features and COSMIC SBS mutational signatures (version 3).

As MuAt features recapitulated mutational patterns reported in literature, we quantified the independent association of each feature with mutational signatures [14, 15]. We found many features to share similarity with single-base (SBS) and doublet-base substitution (DBS), and indel (ID) signatures (Fig. 5; Supplementary Fig. 13 & 14, Supplementary Table 8). These signatures included SBS2 and SBS13 characterized by APOBEC activity (M_15_), SBS17a/b possibly due to DNA damage due to reactive oxygen species or 5FU chemotherapy (M_2_, M_6_, M_22_, M_1_, M_14_), SBS4 due to tobacco smoke (M_3_, M_9_), and SBS12 with currently unknown etiology present in many liver cancers (M_9_). MuAt features explained over half of variance in twelve signatures (Supplementary Fig. 15). Moreover, MuAt features represented patterns across different classes of mutations. For instance, in addition to SBS17, feature M_6_ associated with signature ID2 characterized with single-base deletions at A:T microsatellites attributed to slippage of template strand during DNA replication and observed in large quantities in microsatellite unstable cancers, whereas feature M_21_ associated with signatures SBS9 and ID2, the former attributed to polymerase *η* somatic hypermutation and the latter to slippage during DNA replication of the template DNA strand.

### Attention mechanism captures tumour type specific mutational patterns

To shed light on the mutational patterns learnt by MuAt, we analyzed the similarity matrices *QK*′ extracted from the attention module for the tumours in the PCAWG dataset. Figure 6 shows mutation sequence contexts (“motifs”) for mutation pairs which have received most attention stratified by tumour type. The mutation pair motifs most attended to by MuAt contained a SNV paired either with a SNV (52%), MNV (16%), indel (8%), SV (23%) or MEI (0.4%) (Supplementary Table 9). All these highly-attended MEI events were L1 retrotranspositions occurring in esophageal, pancreatic and prostate adenocarcinomas. Two groups of motifs occurred in most tumour types (Fig. 6 groups A & B). Group A consisted of pairs of SNVs (*e*.*g*., (Tp[C>A]pA, Tp[T>G]A)), group B consisting of SNVs paired with any mutation type (*e*.*g*., (Tp[C>A]pA, Ap[BND]pG), where BND denotes a translocation breakend). In addition to these ubiquitous motifs, many tumour types including PanNETs, brain tumours, and breast, kidney, prostate and thyroid cancers displayed motifs which were characteristic to each tumour type. Well-known mutational patterns appeared among these motifs, such as the doublet C>T substitutions in skin melanomas (Fig. 6 C). We also investigated occurrence of genomic positions in attention matrices. In chronic lymphocytic leukemias and non-Hodgkin lymphomas, attention focused on mutations occurring in the IGH region (Supplementary Fig. 17). These two tumour types displayed both shared and distinct sets of motifs (Fig. 6 D & E). Similarly to motif pair patterns, many tumour types displayed characteristic positional patterns (Supplementary Fig. 17).

**Figure 6:**
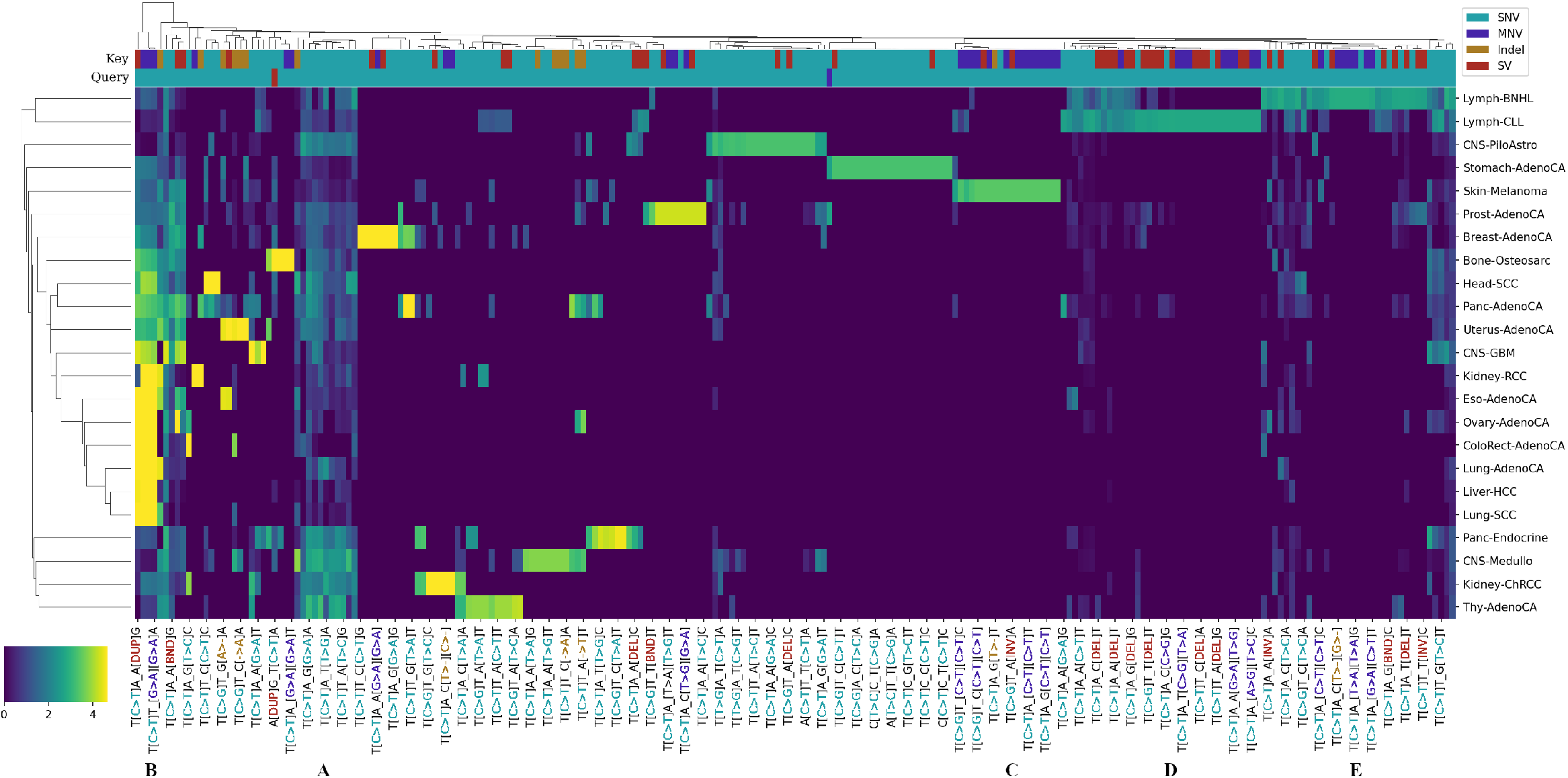
Association between MuAt-derived mutation motifs and tumour types. Attention values for mutation motif pairs (X-axis) extracted from the MuAt model trained on PCAWG data. Values have been averaged first over tumours and then over tumour types (Y-axis). Types of the key and query mutations (SNV, MNV, indel or SV) are indicated on the two top rows. Every fourth motif pair is labelled as “X_Y” where X and Y are the motifs corresponding to attention query and key, respectively. Label colours indicate mutation types. Motif groups: **A**) SNV/SNV and **B**) SNV/non-SNV pairs appearing in many tumour types. **C**) Motifs with doublet C>T substitutions specific to skin melanomas. **D**) and **E**) Motifs characteristic to chronic lymphocytic leukemias and Non-Hodgkin lymphomas.

## 5 Discussion

We introduced a deep neural network called Mutation-Attention (MuAt) to predict tumour types from somatic mutation catalogues while learning representations which are informative of tumour subtypes. To our knowledge, MuAt is the first machine learning model which is able to incorporate heterogeneous information such as mutation type, genomic position and arbitrary annotations on individual mutations instead of representing mutations as aggregated counts. Trained with cancer genomes from the PCAWG consortium, MuAt achieved 89% accuracy in predicting tumour types (24 types), and 74–100% accuracy in independent cohort of cancer genomes. As exome sequencing can be highly informative in clinical settings [46, 47], we trained and evaluated a MuAt model with cancer exomes from the TCGA consortium, and achieved a top-1 accuracy of 64% accuracy. However, top-5 performance in WES data was substantially better, reaching 91% accuracy. The top predictions often included similar tumour types such as lung adenocarcinomas and squamous cell carcinoma, or gastrointestinal tumours. Hence, MuAt results may be informative of tumour origins even if the prediction is not correct. We also observed relatively good performance with downsampled WGS data, suggesting MuAt may find use in low-coverage WGS data, pediatric tumours, and in cell-free DNA applications where only a fraction of the somatic mutation catalogue of a tumour might be captured [22].

We showed MuAt tumour-level features to distinguish between tumour subtypes, even if these labels were not available during training. By associating MuAt features with driver events identified in PCAWG, MuAt highlighted prostate cancers driven by *SPOP* mutations [48], characterised by a 2.3-fold increase in somatic SV burden, exceeding the burden of tumours with *BRCA1* and *BRCA2* mutations. *SPOP* has a role in DNA damage response [49], and *SPOP* mutated prostate cancers have elevated levels of genomic instability [48, 39]. *SPOP* mutations have been associated with better response to therapies [39, 49], potentially mediated by the increased SV burden.

MuAt stratified medulloblastomas into four clusters, one of which contained seven tumours driven by enhancer hijacking events involving *PRDM6* with a 3.7-fold increase in SV burden. *PRDM6* is activated by enhancer hijacking events in Group 4 medulloblastomas [41]. Whether the frequent structural aberrations in these tumours are due to the *PRDM6* activation, or vice versa, remains to be explored in detail. MuAt was also able to highlight a group of four pancreatic neuroendocrine tumours clustering unexpectedly together with kidney cancers. These tumours were found to harbor germline mutations of *MUTYH* with loss-of-heterozygosity in the tumours, explaining the observed clustering [44].

Since MuAt was able to accurately perform tumour typing with somatic mutation catalogues, we hypothesised that the learnt tumour-level features would share a degree of similarity with mutational signatures [14, 15]. We indeed found many MuAt features to associate with COSMIC signatures such as SBS2 and SBS13 (corresponding to APOBEC activity), SBS4 (tobacco smoke), and SBS9 (somatic hypermutation in B cells), although the signatures could not be mapped one-to-one with the features. This is likely due to MuAt not designed to find features which would correspond to independent latent determinants of mutational catalogues similarly to non-negative matrix factorization used in mutational signature methods [14, 15], but instead to predict tumour types as accurately as possible. In contrast to signature analyses requiring a non-trivial refitting step [50, 51], MuAt features for a new sample can be obtained directly given a trained MuAt model. Approaches to learn disentangled representations in deep neural networks may prove fruitful future direction in creating more broadly applicable representations of multiomics data [52, 53].

MuAt’s attention mechanism allowed us to discover aspects of mutation data such as the type and position – or combinations of these – which were informative in predicting each tumour type. We found various types of mutations occurring in the IGH locus to be driving predictions of B cell malignancies. Here MuAt was able to capture the interaction between specific mutation types and the genomic region characteristic to somatic hypermutation in B cells. MuAt was also able to leverage the rarer mutation types such as L1 retrotranspositions to help identify cancers such as esophageal and other epithelial cancers where these events are relatively common [54, 55, 56]. While we were restricted to maximum of 5,000 mutations per tumour due to the complexity *O*(*n*^2^) of the attention mechanism used, recent improvements such as Reformer [57] or Linformer [58] may be used to lift this restriction and to increase MuAt capacity, potentially leading to better performance.

We have demonstrated how deep representation learning can in large cancer datasets yield features which are useful beyond labelled data, for instance in tumour subtyping. Our method, MuAt, can be extended to incorporate additional data on somatic mutations, such as epigenetics, potentially enabling scrutiny of the role of epigenetic interactions in somatic mutagenesis [59, 60, 61, 13, 62]. MuAt is already able to contribute to multiomics data integration to drive biological discovery and clinical applications by providing informative representations of somatic mutation catalogues of tumours. Beyond tumour typing and subtyping, we envision machine learning models such as MuAt to be instrumental in determining cancer prognosis and appropriate treatment choice. As high-throughput patient data accumulates in clinics and cancer projects worldwide, machine learning models able to leverage the massive-scale data will become irreplaceable tools driving digital precision cancer medicine.

## 6 Acknowledgements

This study was supported by grants from the Academy of Finland (322675, 328890), Sigrid Jusélius Foundation, the Cancer Society of Finland, Paulo Foundation, the EI3POD Fellowship programme, the Research Council of Norway (187615), the South-Eastern Norway Regional Health Authority and the University of Oslo. We would like to thank CSC – IT Center for Science, Finland, for generous computational resources. We thank Riku Katainen, Liisa Kauppi, Outi Kilpivaara, Venla Kinanen, Anna Kuosmanen, Laura Langohr and Jaakko Lehtinen for their constructive comments, and Olle Hansson, Jarno Laitinen and Timo Miettinen for technical assistance.

## 7 Methods

### Data acquisition

ICGC and TCGA datasets including consensus somatic variant callsets and sample metadata were obtained from the ICGC data portal (https://dcc.icgc.org/releases/PCAWG/) and Genomic Data Commons data portal (https://portal.gdc.cancer.gov/). All coordinates of somatic variants were specified in GRCh37 human reference genome coordinates.

### Data preprocessing

#### Preparing MuAt inputs from somatic variant callsets

For each somatic variant call in the datasets, MuAt input consists of: 1) mutational sequence (“motif”) encoding for the reference and alternate nucleotide sequence, 2) genomic position (chromosome and position), and 3) additional information on the mutation.

Mutational sequences *s* ∈ Σ^3^ are drawn from alphabet Σ = {A, C, G, T} ∪ ℳ. Mutation symbols ℳ consist of six substitutions described with respect to the pyrimidine base (*i*.*e*., C:G>A:T, C:G>G:C, C:G>T:A, T:A>A:T, T:A>C:G and T:A>G:C), deletions of A, C, G and T, insertions of A, C, G, T, breakpoints for four types of structural variants (*i*.*e*., deletions, duplications, inversions and translocations), and retrotransposon insertions (*i*.*e*., L1, Alu and SINE-VNTR-Alus (SVA)). This encoding allows representing both simple and complex mutations. For instance, the substitution ApCpG>ApTpG would be encoded as A[C>T]G, a diadenine deletion preceded by a cytosine as C[del A][del A], and a deletion breakpoint with a C>G substitution followed by a thymine as [SV_del][C>G]T, where {[C>T], [del A], [SV_del]} ⊂ ℳ.

In experiments, we also considered non-mutation events as negative examples. These are constructed by randomly selecting positions where there is a mutation in one tumour (*e*.*g*., A[C>T]G at chr1:11,235,813), encoding this position without mutational symbols (*e*.*g*., ACG at chr1:11,235,813), and placing it into another tumour’s mutational catalogue. When injecting negative examples into a tumour, we add the median number of variants per type in the dataset as negatives. Specifically, if the dataset has a median of 1000 SNVs and 100 indels, then each tumour will receive a total of 1100 negative examples, where 1000 were picked from random SNVs in other tumours, and the remainder from random indels.

Genomic positions are represented in 1-Mbp bins. For instance, the position (chr1, 11,235,813) would be encoded as the token “chr1_11”. This token is used to encode all mutations occurring in chr:11,000,000–11,999,999.

We experimented with mutational annotations consisting of indicators whether the mutation occurs in a gene (“genic”) or in an exon (“exonic”). In addition, we categorize each mutation into one of four mutually exclusive classes (“strand”): mutation’s pyrimidine reference base is on the 1) same or 2) opposite strand as a gene, or 3) mutation overlaps two genes on opposite strands, or 4) mutation is intergenic. This annotation attempted to capture transcriptional strand biases associated with some mutational mechanisms [63].

Each of the three input modalities (mutation motif, position, annotations) were one-hot encoded separately using token dictionaries. The dictionary of positions consisted 2,915 tokens for all 1-Mb genomic bins. Mutational motif dictionary consisted of 3,692 tokens including 96 SNVs, 2,170 MNVs, 1,160 indels, 233 SVs and 33 MEIs. Finally, the annotation dictionary contained 2×2×4=16 values for the possible combinations of genic, exonic and strand attributes.

### MuAt architecture

The architecture of MuAt is shown in Figure 1, and in detail in Supplementary Figure 1. Input to MuAt is a set of *l* genomic variants. These variants are described with respect to their mutation type and sequence context (mutation motif), genomic position and mutational annotations. Each modality is one-hot encoded using a respective dictionary, resulting in *l* × *m, l* × *p*, and *l* × *g* one-hot matrices, where *m, p* and *g* are the numbers of unique motifs, genomic positions per 1-Mbp bins, and mutation annotations, respectively. These one-hot encoded matrices are then multiplied with embedding matrices ({*m, p, g*} × *k*), resulting in three *l* × *k* embedding matrices. Embedding matrices are concatenated to obtain an *l* × 3*k* matrix *X*_*E*_, which is the input to the MuAt attention module.

Query, key and value matrices are computed by multiplying the input embedding matrix *X*_*E*_ with respective 3*k* × 3*k* weight matrix, *Q* = *X*_*E*_*W*_*Q*_, *K* = *X*_*E*_*W*_*K*_ and *V* = *X*_*E*_*W*_*V*_. The attention mechanism Eq. 1 is then applied *h* times, where *h* is the number of attention heads. The resulting *l* × 3*k* feature matrix is then combined with *X*_*E*_ via skip connections, and fed to batch normalization and fully connected (FC) layers. Finally, the *l* ×3*k* matrix is average-pooled into a 3*k*-vector and processed in a fully connected layer to yield *f* sample-level features (Fig. 4). In our experiments, we used *f* = 24. To obtain the final tumor type predictions, sample-level features are inputted to a fully connected layer, and its outputs are normalized with softmax 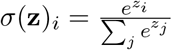 to obtain probabilities over tumor types.

### Experimental design and reporting

We were interested in 1) evaluating the contribution of different mutation types in prediction performance, 2) finding hyperparameters which result in best prediction performance, 3) comparing MuAt with existing models, 4) how to best interpret the trained MuAt models, and whether the features learnt by MuAt are compatible with previous findings. This section provides details how the experiments to answer these questions were prepared.

We report the performance in terms of accuracy *TP* + *TN*/(*TP* + *TN* + *FP* + *FN*), precision *TP/*(*TP* + *FP*), recall *TP*/(*TP* + *FN*) and F1 score 2*TP*/(2*TP* + *FP* + *FN*), where *TP, TN, FP, FN* are the number of true positives and negatives, and false positives and negatives, respectively. Top-*k* accuracies were calculated such that the prediction was deemed correct when the correct class is among the *k* highest scoring predictions.

### MuAt hyperparameter search and model training

We performed search for MuAt hyperparameters over embedding dimensions {128, 256, 512}, number of encoder layers {1, 2, 4}, number of attention heads {1, 2}, number of fully connected layers {1, 2} and mutation types to be included in the input. Supplementary Figure 2 shows the mutation type combinations for PCAWG (15 combinations) and TCGA datasets (9 combinations). We set the learning rate to 6 × 10^−4^, momentum 0.9, and minibatch size of one, training for 150 epochs. Maximum number of mutations MuAt was able to process per tumour in our experiments was 5000, limited by the memory on GPUs available to us (see Programming environment). MuAt parameters were optimized with stochastic gradient descent minimizing cross-entropy loss

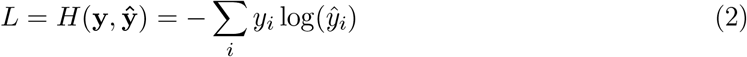

where *ŷ*_*i*_ is predicted probability for tumour type *i*, and *y*_*i*_ ∈ {0, 1} denotes whether *i* is the correct tumour type.

We performed 10-fold cross-validation by splitting datasets into training and validation sets with the ratio of 9:1 to train MuAt models. In each fold, we trained a model for each combination of the hyperparameter choices listed above, and the model achieving the best accuracy in the validation set was selected from each fold. The hyperparameters and prediction performance for each fold is given in Supplementary Table 3. The model selected for analysis of tumour typing performance in sparse data and tumour subtype discovery was trained on SNVs, MNVs, indels and SVs/MEIs, genomic positions and mutation annotations using the PCAWG dataset. This model contained one attention head and two encoder layers, embedding dimension 512, resulting in 28,458,520 trainable parameters.

### Comparing MuAt with other models

We compared MuAt to the deep neural network proposed by Jiao *et al*. [20] and a random forest (RF) model. We used the default setting of RF regressor available in scikit-learn package.

In [20], a total of 150 SNV features were used (SNV150) containing the six possible singlenucleotide substitutions (C>A, C>G, C>T, T>A, T>C, T>G), SNVs together with either the flanking 5’ or 3’ base (4×6 + 6×4 = 48 features), and SNVs together with both the 5’ and 3’ flanking bases (4×6×4 = 96 features).

We performed 10-fold cross-validation to train the DNN model. DNN model hyperparameters were optimized in each fold as done in [20]. For the random forest model, we performed 4-fold cross-validation.

The contribution of attention mechanism to MuAt performance was evaluated by comparing a MuAt model with one encoder layer and one attention head to a model without any encoder layers.

#### Evaluating MuAt in sparse and independent data

To test the performance of MuAt in sparse data, and further to test transfer learning from WGS to WES data. We selected 13 common tumour types existing in both PCAWG and TCGA dataset (Supplementary Table 1). With the same hyperparameters setup as mentioned previously, we retrain the model, as well downsampled validation set with fixed number of mutations *n* ∈ {10, 50, 100, 300, 1000, 2500, 4000, 5000}, where 5000 is the maximum capacity of MuAt. Results for transfer learning are shown in Supplementary Figure 6. This setup also used for testing the effect of attention (Fig. 3 and Supplementary Fig. 5)

To evaluate MuAt in independent data 7, we created an ensemble MuAt model by taking the ten best performing, cross-validated models trained on PCAWG data, and predicting tumor type based on summed logits for tumor types from each model.

#### Extracting and visualising MuAt tumour-level features

To visualize the tumour-level features learnt by MuAt, we extracted the output of the layer before the final prediction (Supplementary Fig. 1). This yielded a feature vector of length 24 for each tumour, which we projected onto a two-dimensional space with UMAP [36]. Interactive UMAP visualization is available at https://github.com/primasanjaya/mutation-attention.

#### Association of MuAt features with driver events and somatic mutation patterns

Driver events identified in PCAWG tumours [33] were associated with principal components of MuAt features with least-squares regression. For each pair of MuAt feature principal component (*n*=24) and driver (*n*=298), a least-squares linear model *p* ∼ *d*+age+sex+histology+*g*_1_+…+*g*_10_ was fitted, where *p* is the feature principal component, *d* is an indicator whether the driver event was detected in the tumour, and *g*_*i*_ is the *i*th principal component of patient genotypes computed in PCAWG [33]. Association was computed only for drivers with at least three tumours harboring the driver, resulting in 7,128 models. *P* -values from all tests were adjusted for multiple testing with Benjamini-Hochberg method. Supplementary Figure 8 shows the driver coefficients for all feature principal components from all models, as well as a histogram and a quantile-quantile plot of unadjusted p-values against a uniform distribution showing relatively small degree of inflation.

To analyse correspondence of MuAt tumour-level features with COSMIC signatures, we first calculated Spearman’s rank correlation between each MuAt feature (*n*=24) and COSMIC SBS signature (Fig. 5). We also carried out least-squares linear regression for each signature separately to predict the log-transformed signature value *s* based on all MuAt features, *i*.*e*., log(*s*) ∼ *M*_1_ + … + *M*_24_. We corrected *p*-values with the Benjamini-Hochberg method [64] and reported results with false discovery rate (FDR) <10% (Supplementary Fig. 13, 14). For each signature, the variance explained by MuAt features as adjusted *R*^2^ value is given. This analysis was performed for both COSMIC version 2 and version 3 signatures.

Association of MuAt features with mutation counts stratified by type was quantified with negative binomial model to predict mutation count based on MuAt features *M*_1_, …, *M*_24_, showing results with FDR<5% in Supplementary Figure 16. Association with MSI levels was performed by predicting the logarithm of fraction of mutated microsatellites computed previously [65] from MuAt features, *i*.*e*., MSI ∼ M_1_, …, M_24_ with a least-squares regression model.

#### Inspecting attention matrices

We analysed the attention matrices *QK*′ of each tumour in the PCAWG data by first extracting the 5000×5000 matrices *A* = (*a*_*ij*_), and selecting the values *a*_*ij*_ > 0.9×max(*A*) to reduce the size of data. Rows and columns of *A* correspond to mutations of a tumour. We can thus visualise the matrices with respect to different mutational data modalities; in our experiments, we visualised mutational motifs and genomic positions. Genomic annotations, (*i*.*e*., genic, exonic and strand attributes) were not visualised.

To create Figure 6 and Supplementary Figures 17, we first expressed the attention values in terms of the selected modality (*e*.*g*., mutation motifs for query and key mutations in Fig. 6), then averaged the values over tumours and tumour types, and finally divided by column(modalities) and row-wise (tumour types) standard deviations.

#### Programming environment

We implemented MuAt with PyTorch 1.8.0 deep learning framework in Python 3.7. To evaluate the model of Jiao *et al*. [20], we used the code provided at https://github.com/ICGC-TCGA-PanCancer/ TumorType-WGS, and ran it with TensorFlow 2.0 in Python 3.6. For the random forest model, we used scikit-learn 0.21.3. Statistical modelling was done with scipy 1.5.3 and statsmodels 0.12.1 packages. Packages used for data analysis and visualization included pandas 1.3.4, seaborn 0.11.2 and umap-learn 0.5.1. All deep neural network models were trained on NVidia Tesla V100 GPUs with 16 GB memory.

