## Supplementary Figures 1-17 for "Mutation-Attention (MuAt): deep representation learning of somatic mutations for tumour typing and subtyping"

[3]iCAN Digital Precision Cancer Medicine Flagship, Finland

[4]Centre for Molecular Medicine Norway (NCMM), Nordic EMBL Partnership, University of Oslo and Oslo University Hospital, Oslo, Norway

[5]Department of Pediatric Research, Division of Pediatric and Adolescent Medicine, Rikshospitalet, Oslo University Hospital, Oslo, Norway

[6]Department of Neurology, University of California, San Francisco, San Francisco, United States

[7]Division of Computational Genomics and Systems Genetics, German Cancer Research Center (DKFZ), Heidelberg, Germany

[8]Genome Biology Unit, European Molecular Biology Laboratory, Heidelberg, Germany

[9]European Molecular Biology Laboratory, European Bioinformatics Institute, Wellcome Genome Campus, Hinxton, Cambridge, UK

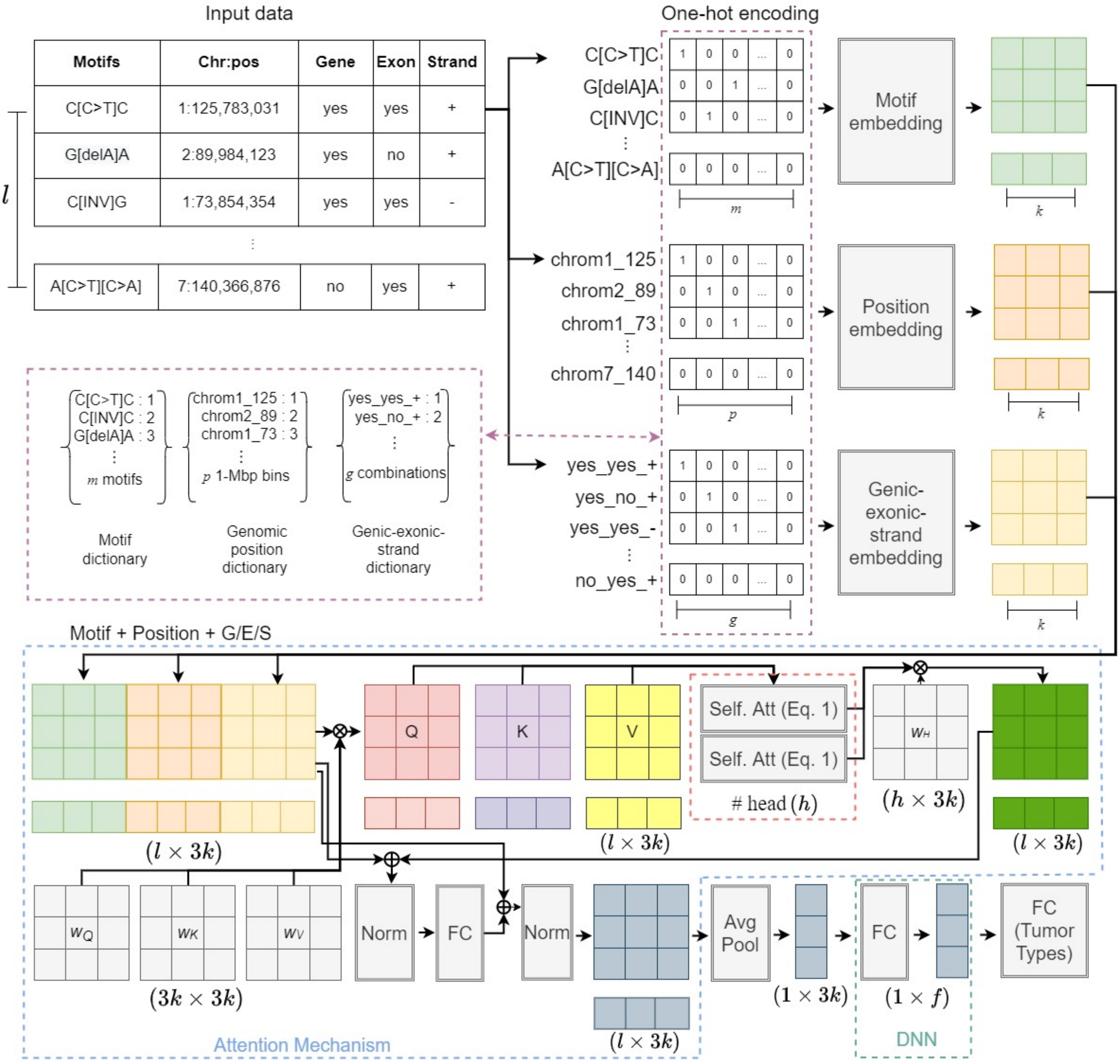


Supplementary Figure 1: Architecture of MuAt.

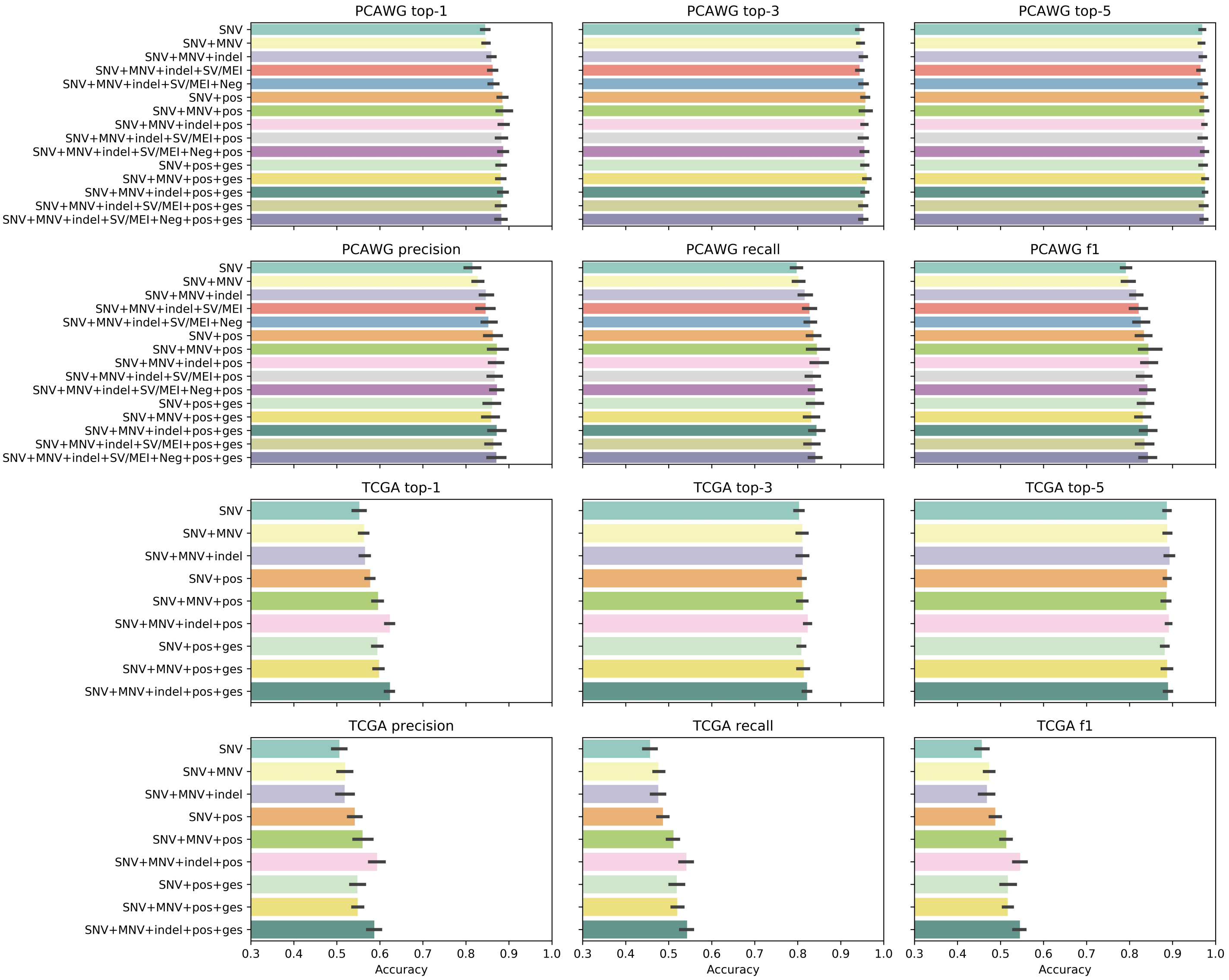


Supplementary Figure 2: Top-1, top-3 and top-5 accuracy, precision, recall and F1 scores of MuAt models trained with different mutation types in PCAWG (top) and TCGA (bottom) data. Tests where negative examples were injected into data (see Methods) are denoted with "Neg". Tests where genic, exonic and strand attributes (Methods) were used are denoted with "ges".

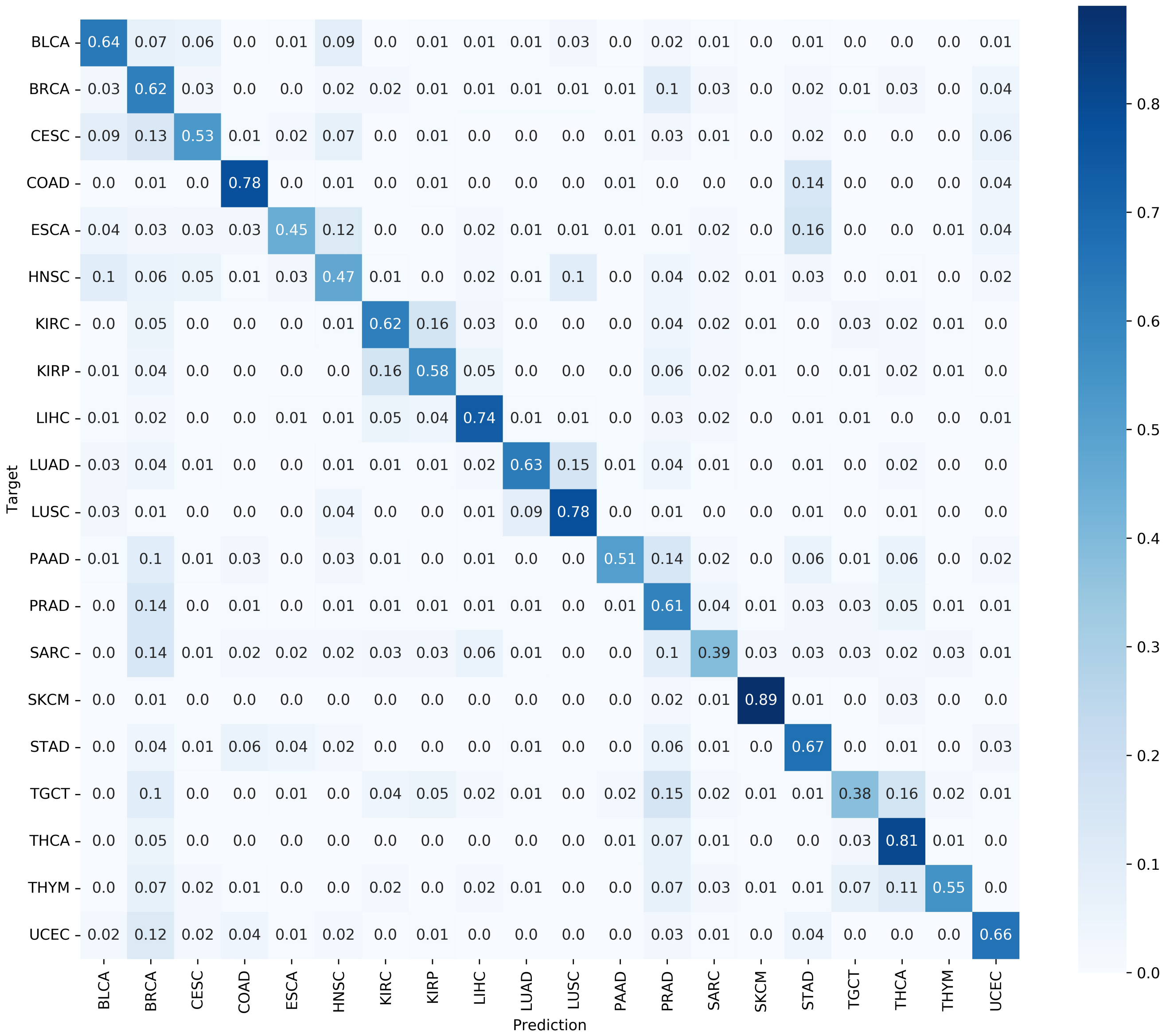


Supplementary Figure 3: Confusion matrix of MuAt in TCGA data.

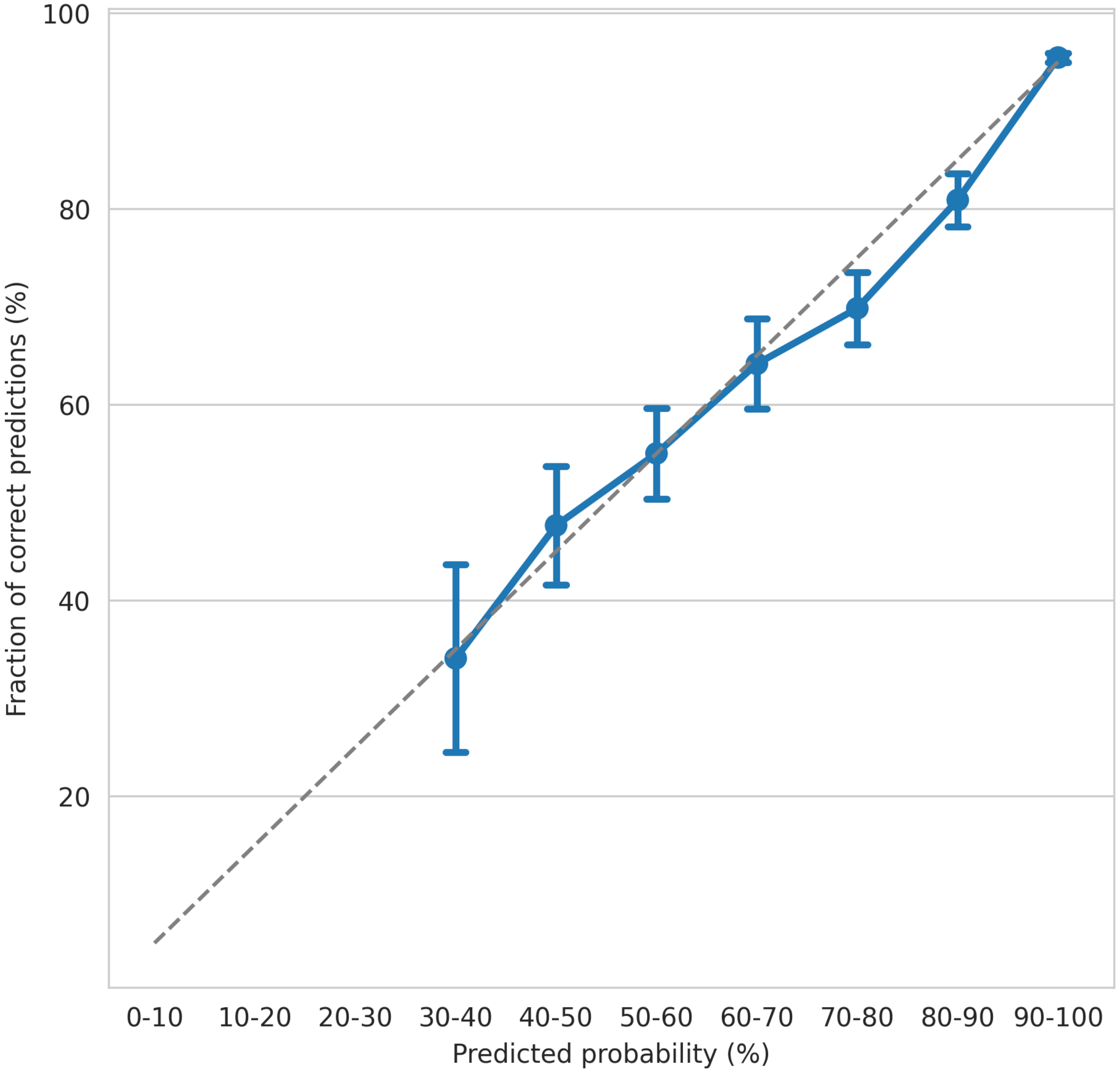


Supplementary Figure 4: Calibration curve of MuAt trained with PCAWG data showing the binned predicted probabilities (X-axis) versus the fraction of positive predictions (Y-axis). Error bars indicate standard deviation in bootstrapped data (n=1000). Only bins with more than three tumours shown, excluding the bin [20%,30%) with three tumours.

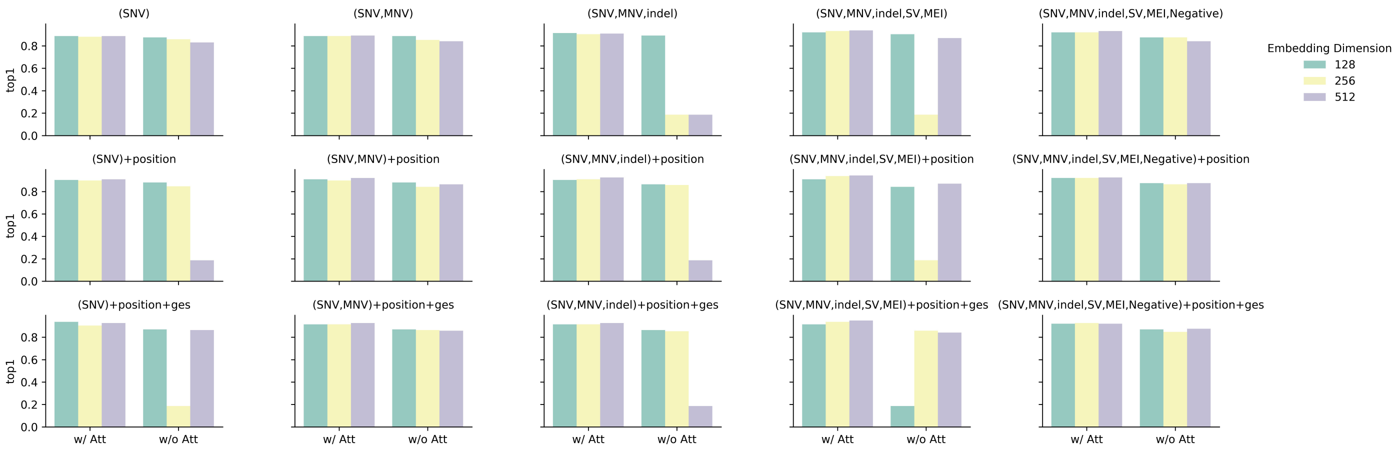


Supplementary Figure 5: MuAt model performance with (w/) and without (w/o) the attention module in PCAWG data with different combinations of mutation types.

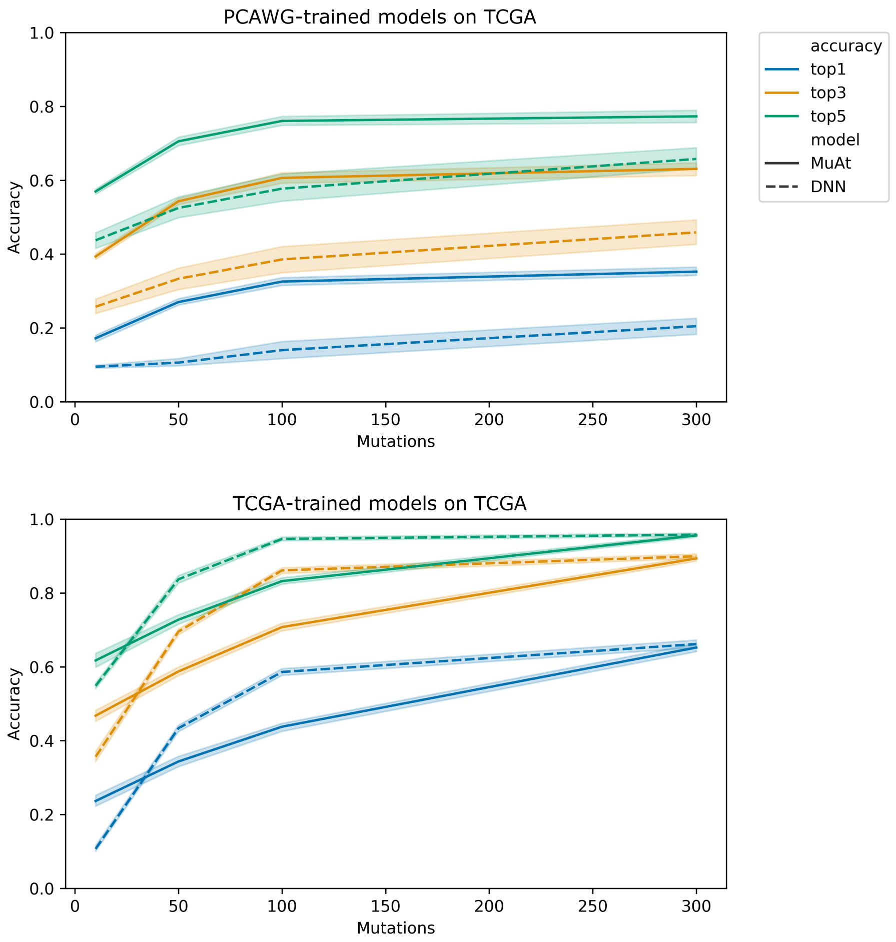


Supplementary Figure 6: Transfer learning performance. Top: PCAWG-MuAt on TCGA data. Bottom: TCGA-MuAt on TCGA data. X-axis: maximum number of mutations sampled from each tumour, Y-axis: prediction accuracy.

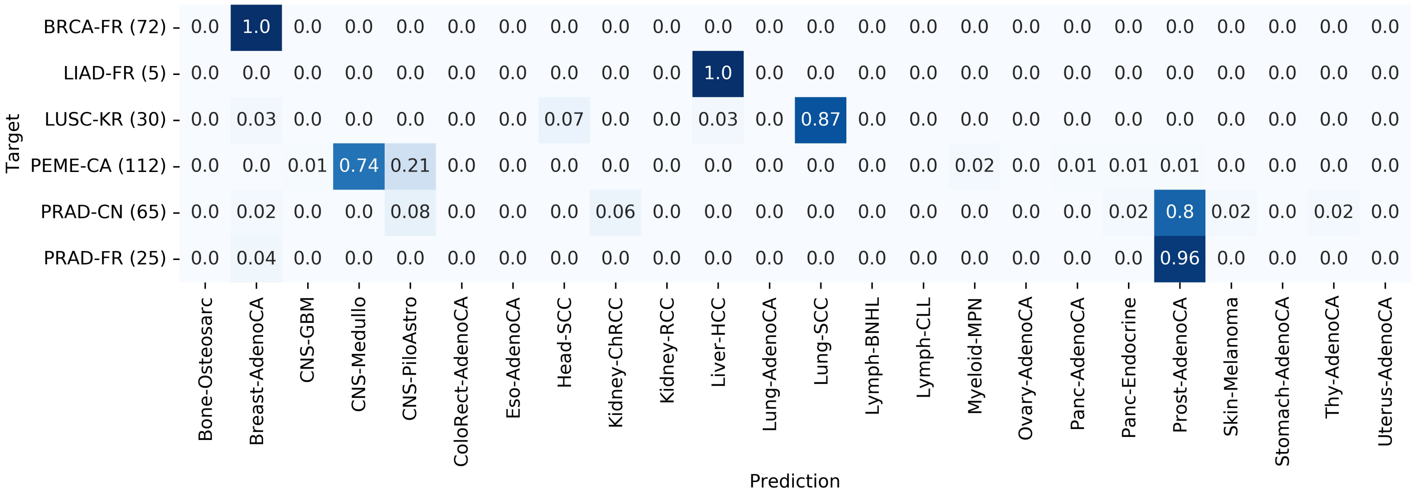


Supplementary Figure 7: Confusion matrix for MuAt models trained on PCAWG SNVs and MNVs with genomic positions and mutation annotations in the independent ICGC dataset. X-axis: tumour types in PCAWG training data, Y-axis: tumour types of the independent ICGC dataset: Breast Cancer (BRCA-FR), Benign Liver Tumour (LIAD-FR), Lung Cancer – Squamous cell carcinoma (LUSC-KR), Pediatric Medulloblastoma (PEME-CA) and Prostate Cancer (PRAD-CN, PRAD-FR).

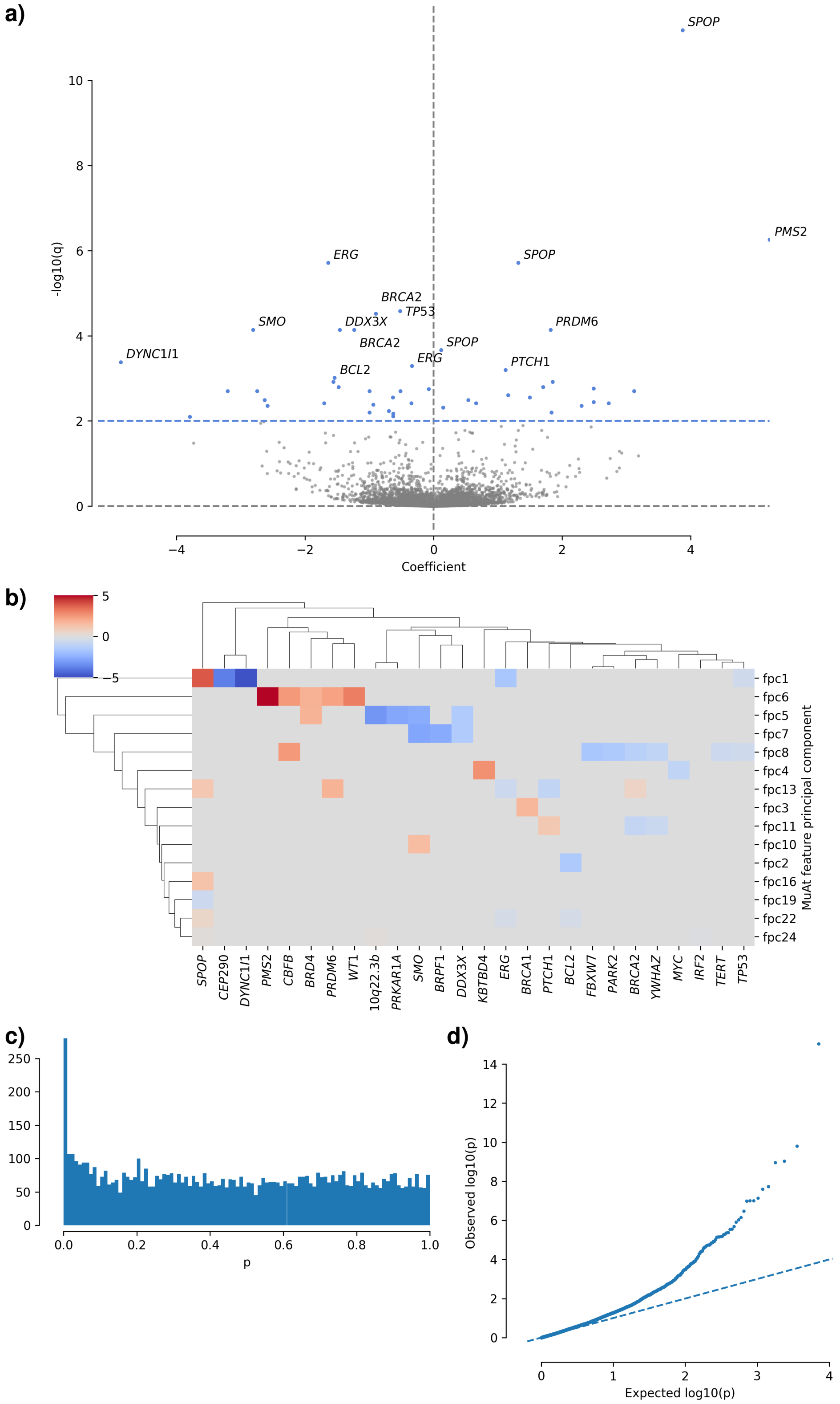


Supplementary Figure 8: Linear association of PCAWG driver events with MuAt tumour-level feature principal components (FPCs). Least-squares regression model was corrected with patient age and sex, tumour histology, and first ten principal components of patient genotype. **a)** Volcano plot showing coefficients of all gene-FPC pairs (X-axis) and $\log_{10}(q)$ values (Y-axis). FDR<1% associations are indicated with blue colour, and top 15 associations are labeled. **b)** A clustered heatmap of association coefficients8 **c)** Histogram and **d)** quantile-quantile plot of $p$-values.

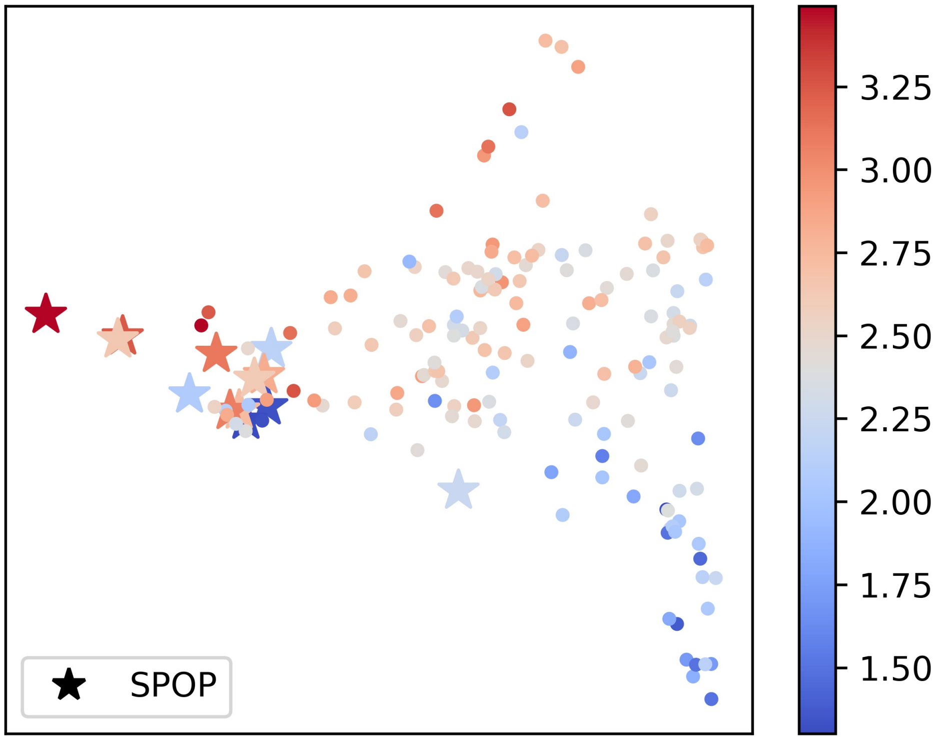


Supplementary Figure 9: Prostate cancers with *SPOP* driver events. X- and Y-axis: UMAP components of tumour-level features in the MuAt PCAWG model (coordinates are the same as in Figure 4). Colouring indicates structural variant burden in a log scale.

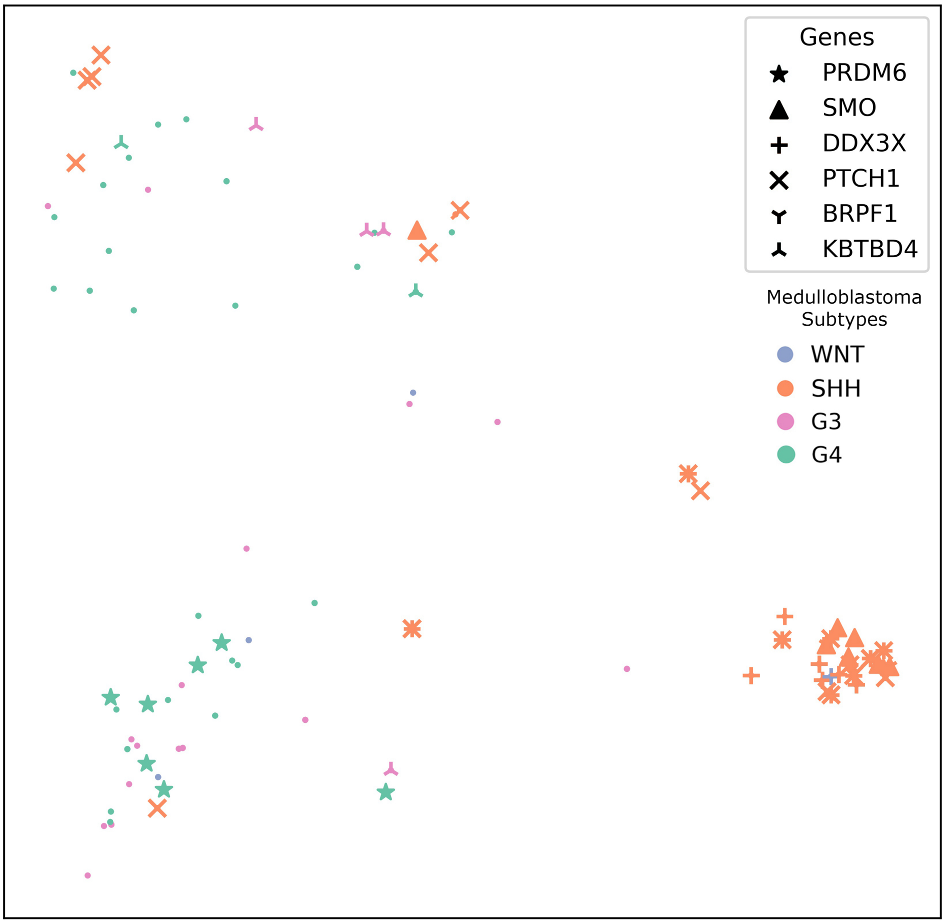


Supplementary Figure 10: MuAt tumour-level feature UMAP showing medulloblastomas in the PCAWG dataset (coordinates are the same as in Figure 4). Tumours with driver events associating with MuAt features are highlighted.

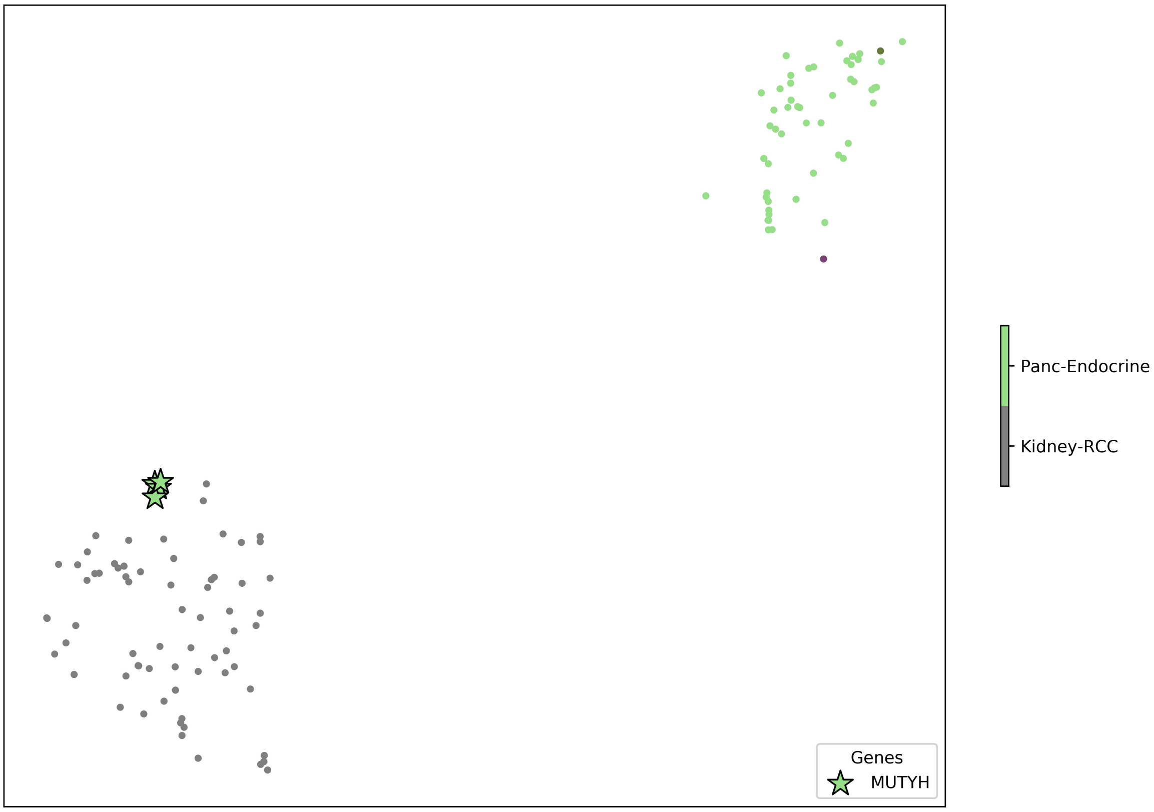


Supplementary Figure 11: Pancreatic endocrine tumours of four patients clustering with kidney cancers in MuAt tumour-level feature UMAP (coordinates are the same as in Figure 4). These four patients carry germline mutations of *MUTYH*, with loss-of-heterozygosity observed in all four tumours.

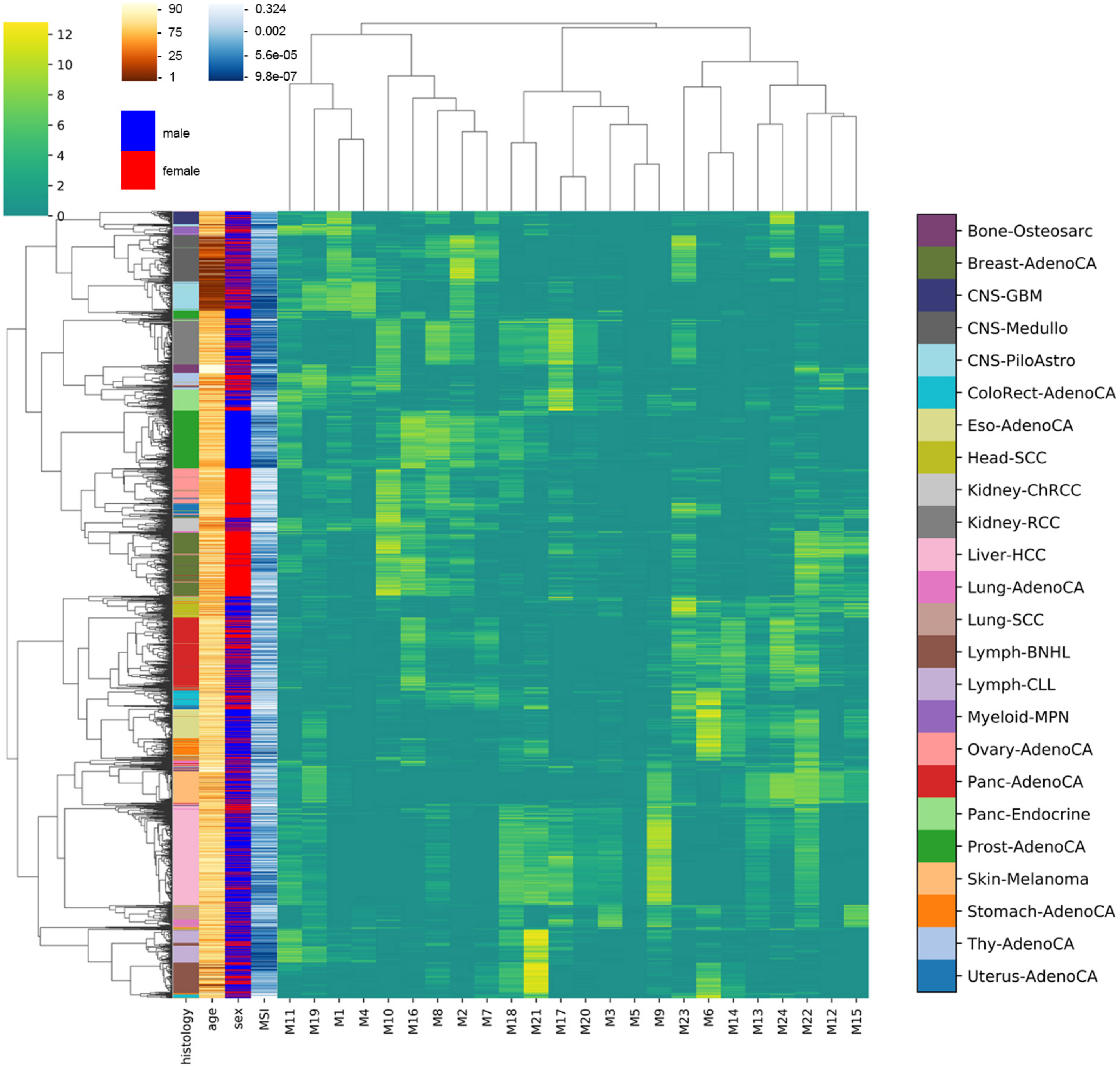


Supplementary Figure 12: A clustered heatmap of MuAt tumour-level feature with age, sex, and MSI values in each PCAWG tumours.

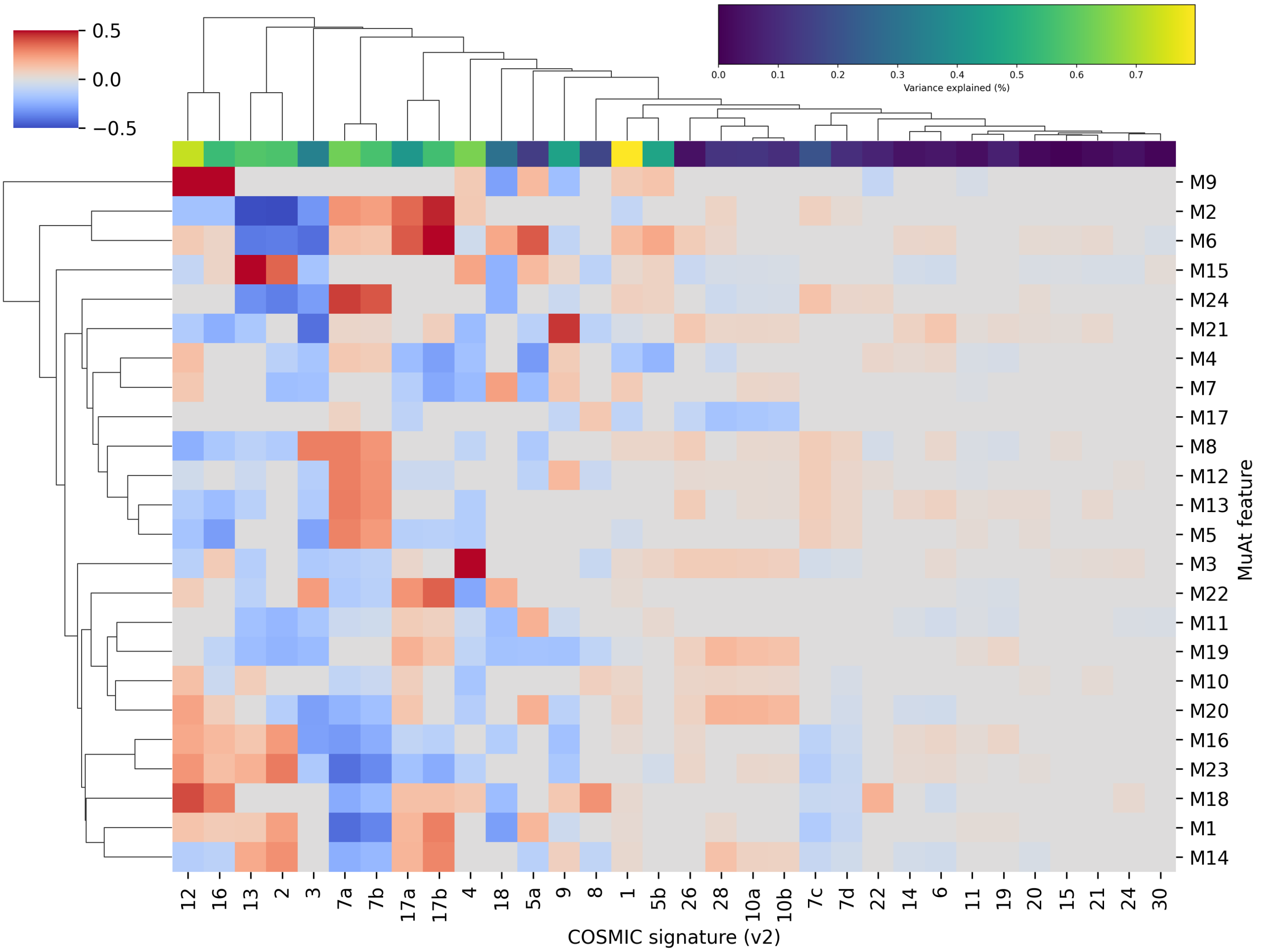


Supplementary Figure 13: Association of MuAt PCAWG model tumour-level features (Y-axis) with COSMIC mutational signatures (version 2; X-axis). Log-transformed number of mutations attributed to each signature was predicted with least-squares regression given MuAt features. FDR<10% associations shown in non-grey colours. Variance of each signature explained indicated on the top row.

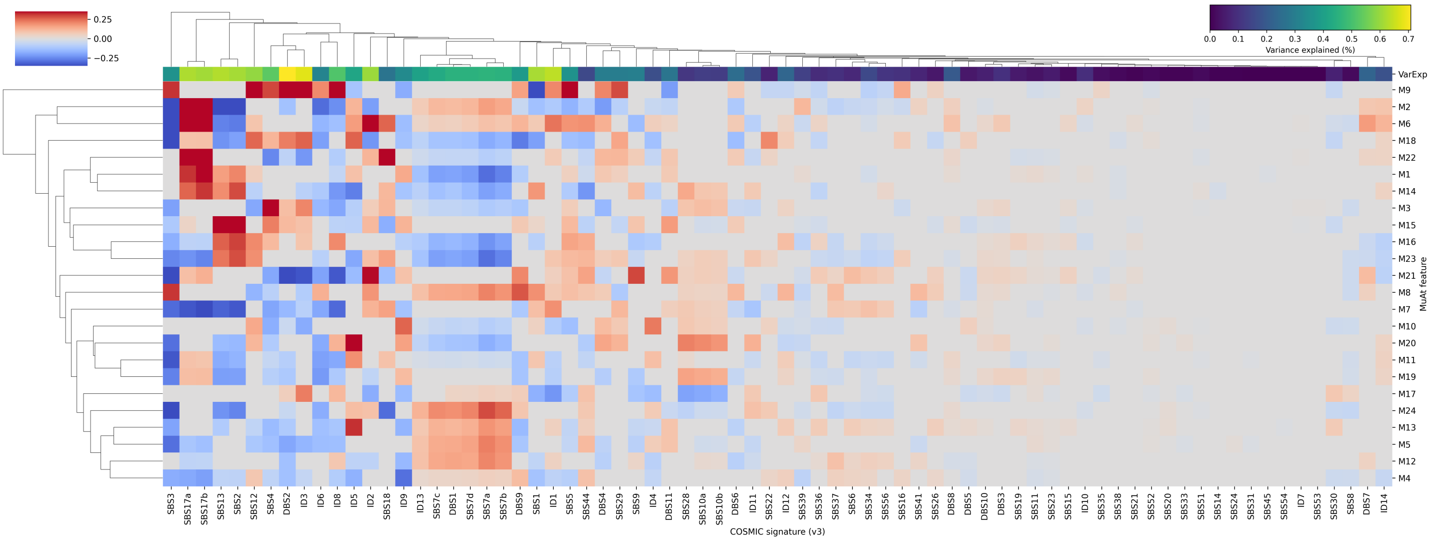


Supplementary Figure 14: Association of MuAt PCAWG model tumour-level features (Y-axis) with COSMIC mutational signatures (version 3; X-axis). Log-transformed number of mutations attributed to each signature was predicted with least-squares regression given MuAt features. FDR<10% associations shown in non-grey colours. Variance of each signature explained by the regression model indicated on the top row (see also Supplementary Fig. 15.

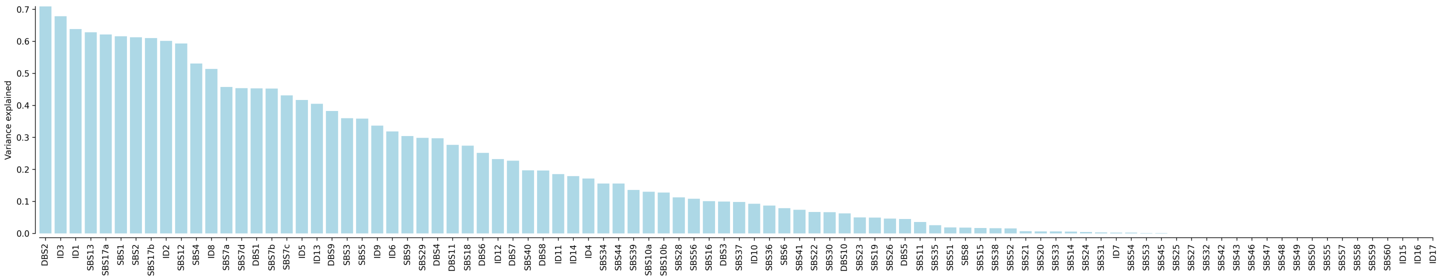


Supplementary Figure 15: Variance in the number of mutations attributed to signatures (COSMIC version 3) explained by MuAt tumour-level features.

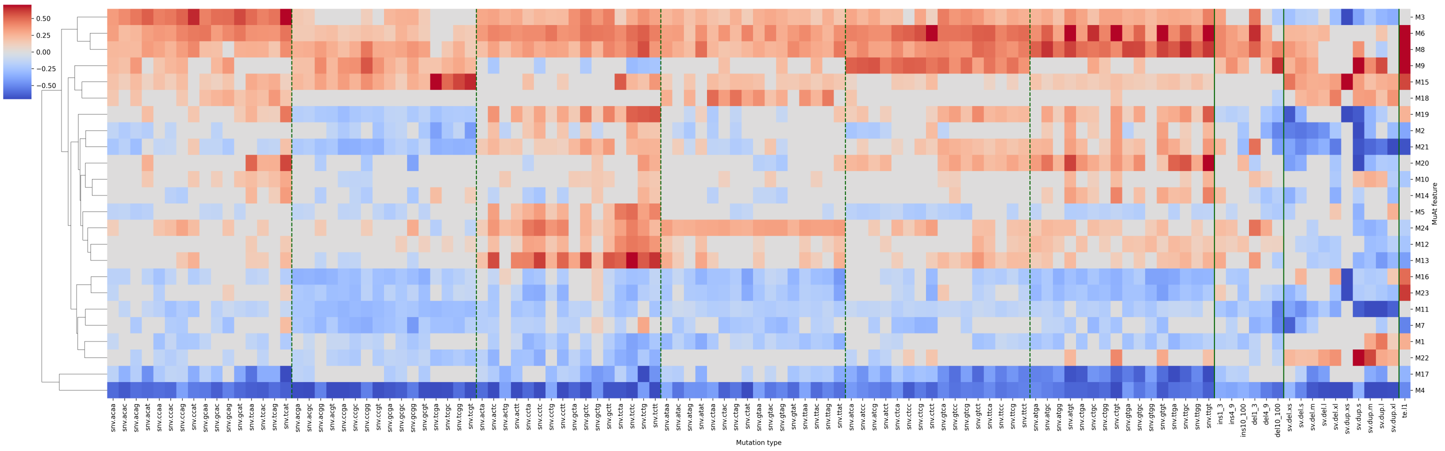


Supplementary Figure 16: Association of MuAt tumour-level features (Y-axis) with mutation counts by type (X-axis). A negative binomial model was used to predict counts of each mutation type given MuAt features M1,. . .,M24 in PCAWG data. FDR<5% results shown as non-grey colours.

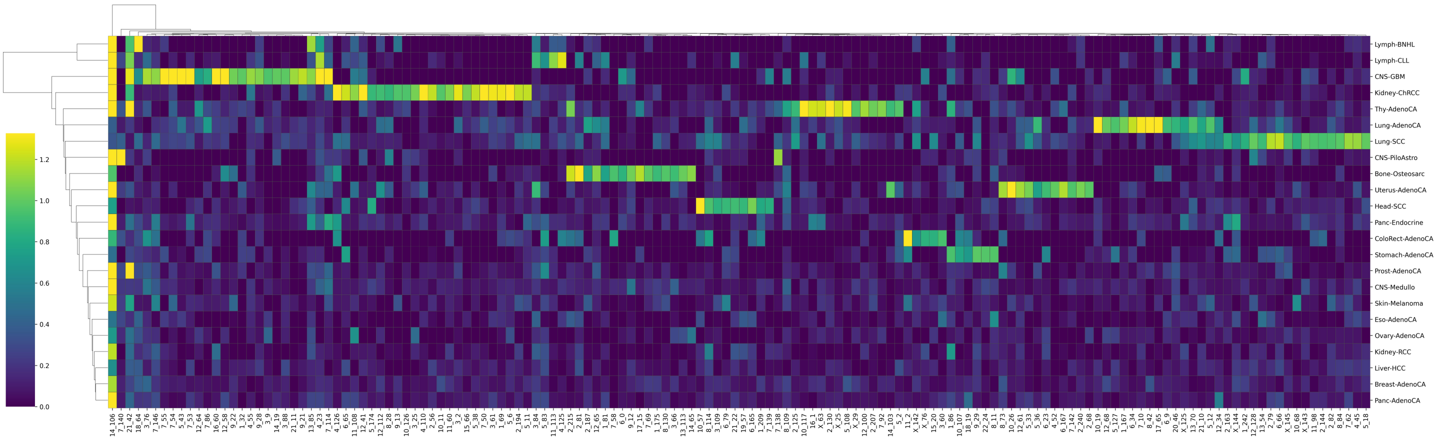


Supplementary Figure 17: Mean attention values of genomic positions (X-axis) per tumour type (Y-axis) in the MuAt PCAWG model. Genomic positions given as "chromosome_$b$", where $b$ is the 1-Mbp bin starting at chromosome position $b \times 10^6$. Genomic positions with the highest 5% variability in attention across tumours are shown. Many genomic position bins appear characteristic to specific tumour types. Genomic region chr14:106,000,000-107,000,000 contains the *IGH* region (leftmost bin), which is recurrently mutated in B cells.
